# Nanocrown electrodes for reliable and robust intracellular recording of cardiomyocytes and cardiotoxicity screening

**DOI:** 10.1101/2021.09.28.462181

**Authors:** Zeinab Jahed, Yang Yang, Ching-Ting Tsai, Ethan P. Foster, Allister F. McGuire, Huaxiao Yang, Aofei Liu, Csaba Forro, Zen Yan, Xin Jiang, Ming-Tao Zhao, Wei Zhang, Xiao Li, Thomas Li, Annalisa Pawlosky, Joseph C. Wu, Bianxiao Cui

## Abstract

Drug-induced cardiotoxicity arises primarily when a compound alters the electrophysiological properties of cardiomyocytes. Features of intracellular action potentials (iAPs) are powerful biomarkers that predict proarrhythmic risks. However, the conventional patch clamp techniques for measuring iAPs are either laborious and low throughput or not suitable for measuring electrically connected cardiomyocytes. In the last decade, a number of vertical nanoelectrodes have been demonstrated to achieve parallel and minimally-invasive iAP recordings. Nanoelectrodes show great promise, but the large variability in success rate, signal strength, and the low throughput of device fabrication have hindered them from being broadly adopted for proarrhythmia drug assessment. In this work, we developed vertically-aligned and semi-hollow nanocrown electrodes that are mechanically robust and made through a scalable fabrication process. Nanocrown electrodes achieve >99% success rates in obtaining intracellular access through electroporation, allowing reliable and simultaneous iAP recordings from up to 57 human pluripotent stem-cell-derived cardiomyocytes (hPSC-CMs). The accuracy of nanocrown electrode recordings is validated by simultaneous patch clamp recording from the same cell. Nanocrown electrodes enable prolonged iAP recording for continual monitoring of the same cells upon the sequential addition of four to five incremental drug doses. In this way, the dose-response data is self-referencing, which avoids the cell-to-cell variations inherent to hPSC-CMs. We are hopeful that this technology development is a step towards establishing an iAP screening assay for preclinical evaluation of drug-induced arrhythmogenicity.

## Introduction

The waveforms of intracellular action potentials (iAP) reflect the coordination of a multitude of ion channels, some of which are affected by drugs to collectively contribute toward proarrhythmic risks. Parameters including the action potential duration, diastolic interval, rate of depolarization, and the triangulation of repolarization of iAPs, have been shown to serve as predictors of proarrhythmia using human pluripotent stem-cell-derived cardiomyocytes (hPSC-CMs) (Hondeghem and Hoffmann 2003; Colatsky et al. 2016). The measurement of iAPs is mostly performed by patch clamp, which compares iAP waveforms before and after multi-dose drug treatment from the same cell, i.e. self-referencing. Manual patch clamp is reliable and accurate, but is limited by low throughput and high labor costs. For example, most manual patch clamp studies measure 3-7 hPSC-CMs per drug dose (Gibson et al. 2014; Liang et al. 2013; Crumb et al. 2016; Hyun et al. 2017). Several automated patch clamp techniques have been shown to increase the throughput (Scheel et al. 2014; Kramer et al. 2020; Dunlop et al. 2008), but these methods require the use of isolated cells in suspension, which disrupts electrically connected cardiomyocytes and may compromise the relevance of the measurement. Even with automated patches, studies usually measure <10 cells per drug dose (Scheel et al. 2014; Li et al. 2019; Stoelzle et al. 2011). Noninvasive and multiplexable microelectrodes can easily achieve parallel recording of extracellular action potentials (eAPs). However, eAPs appear as biphasic spikes and do not provide all crucial information for proarrhythmia assessment. Recent microelectrode-based local extracellular action potential method can capture many iAP features but it lacks one-to-one cell to electrode correspondence, is not self-referencing, and the accuracy of measured iAP features is yet to be fully validated by patch clamp (Hayes et al. 2019; Lopez et al. 2018; Edwards et al. 2018).

In the last decade, vertically-aligned and solid-state nanoelectrode arrays (NEAs) have emerged as promising tools with the potential of achieving parallelizable and minimally invasive iAP recording from monolayers of substrate-adhered cardiomyocytes (Abbott et al. 2018; McGuire, Santoro, and Cui 2018; Robinson et al. 2012; Xie et al. 2012; Duan et al. 2011; Hai, Shappir, and Spira 2010; Angle, Cui, and Melosh 2015; Spira and Hai 2013; B. X. E. Desbiolles et al. 2019; Benoît X. E. Desbiolles et al. 2020). The recent development of combining vertical nanoelectrodes with CMOS technology has drastically increased the recording throughput of NEAs and has achieved high-throughput recordings in both cardiomyocyte cultures and neuronal networks (Abbott et al. 2020, 2017). Additionally, enabled by integrated current clamp electronics, these methods can now reliably perform intracellular AP recording from Neurons which was previously only achievable by patch clamp (Abbott et al. 2020). NEA devices differ substantially in their shapes, throughput, and mechanisms of gaining intracellular access. Some devices appear to spontaneously gain intracellular access (B. X. E. Desbiolles et al. 2019; Hai, Shappir, and Spira 2010; Robinson et al. 2012; Liu et al. 2017, 2021), while others require transient electroporation (Abbott et al. 2017; Lin et al. 2014; Benoît X. E. Desbiolles et al. 2020; Xie et al. 2012; Lin et al. 2017; Abbott et al. 2020) or optoporation (Dipalo et al. 2018, 2021, 2017) of the cell membrane. Although spontaneous intracellular access allows iAP recording, it often occurs randomly with a low probability, and cannot be repeated in a controllable manner (B. X. E. Desbiolles et al. 2019). On the other hand, transient electroporation of the membrane with a short electric pulse is a controlled method of repeatedly gaining intracellular access and recording iAPs at desired time points. Although all NEAs should be compatible with self-referencing recording, the variable success rate and recording duration have largely prevented their usage for multi-dose drug screening. In addition, the fabrication of the majority of nanoelectrodes involve electron beam or ion beam lithography, which are costly and serial processes. These limitations have so far hindered NEAs from being broadly adopted for drug screening purposes.

Other approaches are being developed as surrogate assays for the measurement of iAPs, including voltage and Ca^2+^ optical imaging using fluorescent indicators or genetically encoded molecular probes (Broyles, Robinson, and Daniels 2018; Dana et al. 2019; Shroff et al. 2020). These powerful approaches allow simultaneous recording from many cells. That said, some significant drawbacks still exist for optical approaches including the requirement of chemical labeling or genetic modification of cells, photo-toxicity, limited duration of measurement due to photobleaching, and slower on/off kinetics. All these constraints currently prevent them from replacing direct electrophysiological methods for recording iAPs. Rather, it is often desirable to combine electrophysiological and optical interrogation of the same cell for multimodal analysis, such as correlation between iAPs and calcium transients in cardiomyocytes.

Herein, we developed semi-hollow nanocrown electrodes that are mechanically robust and can perform highly reliably and prolonged iAP recordings. Nanocrown electrodes enable parallel, self-referencing, and multi-dose drug assessment in hPSC-CMs. When combined with a miniaturized recording apparatus housed inside an incubator, these nanoelectrodes will enable iAP-based drug screening.

## Results and Discussion

### Fabrication of robust nanocrown electrode arrays on transparent substrates by photolithography

Our device fabrication process consists of two parts. In the first part of the device fabrication, we developed a protocol to address issues related to the robustness and the fabrication throughput of vertical nanoelectrode arrays (NEAs). In this protocol, photolithography and a top-down dry etching technique were used to fabricate transparent SiO_2_ nanopillars (∼1 µm in diameter and ∼3 µm in height) by directly etching into a SiO_2_ substrate (**Fig. 1A)**. These SiO_2_ nanopillars are mechanically robust as they are continuous with the substrate at their base. The dry-etching chemistry was optimized to obtain a slightly tapered micropillar profile with a larger diameter and stronger attachment near the base. To reduce the feature size to nanoscale (Lou et al. 2019; W. Zhao et al. 2017), we employed a wet SiO_2_ etching technique that linearly and uniformly shrinks vertical pillars from 1000 nm to 500 or even 200 nm diameter (**Fig. 1B**). The final diameter is controlled by the duration of the wet etch step. Subsequently, we used photolithography and Pt deposition to pattern conducting electrodes (surrounding the SiO_2_ nanopillars) and connecting feed lines. The tapered nanopillar favors the metal coverage and continuity at the nanopillar-substrate edges during metal sputtering. Finally, a thick 600-nm insulating layer of SiO_2_/SiN_x_ was deposited to insulate the entire surface.

**Figure 1.**
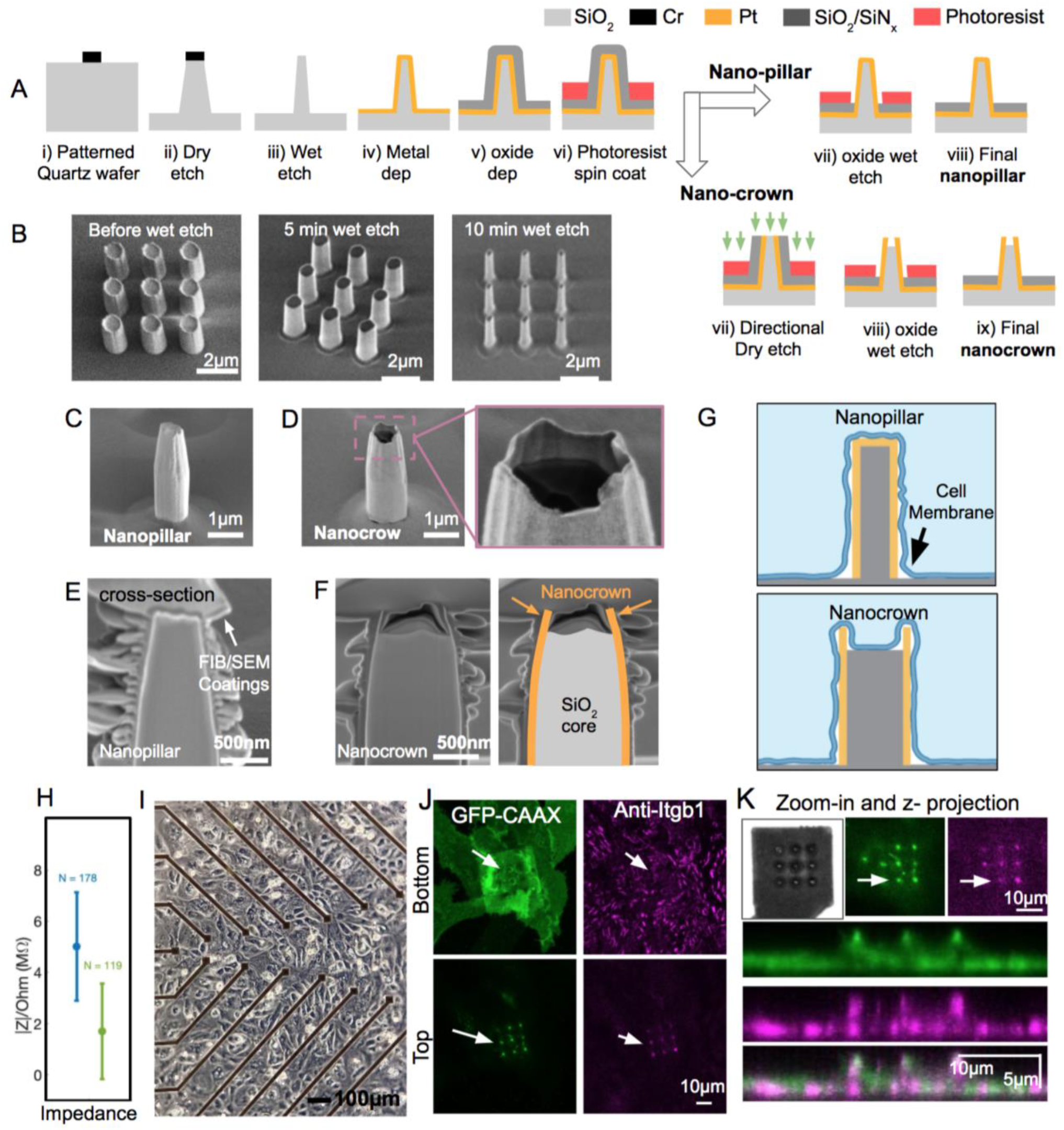
Fabrication and characterization of robust nanopillar and nanocrown electrodes arrays (NEAs) by optical lithography. A) Illustrations of the nanofabrication steps for NEAs. (i) Circular holes were patterned on transparent SiO_2_ wafers using maskless photolithography, and Cr masks were evaporated in the patterned holes. (ii) Reactive ion etching was used to etch vertically into the SiO_2_ substrate and produce slightly-tapered vertical standing pillars of 700 nm diameters. (iii) The pillars were subsequently thinned down to 200 nm diameter using a buffered oxide (BOE) wet etch. (iv) Pt metal was sputter coated onto the nanopillars to produce conductive vertical nanopillars. (v) An SiO_2_/Si_3_N_4_ layer was then evaporated onto the surface for insulation. vi) A photoresist was coated around the nanopillar to protect the insulation layer for subsequent steps. In the next steps, either nanopillar or nanocrown electrodes were fabricated from the nanopillar through different processes. To fabricate nanopillars, an oxide wet etch was used to remove the insulation layer from the pillar tops. To fabricate nanocorwn electrodes, first a directional dry etch was used to remove the SiO_2_/Si_3_N_4_ and Pt metal layers from the pillar tips. Next the SiO_2_ base of the nanopillar was etched down by a controlled oxide wet etch to fabricate nanocrowns with 50-200nm depths. B) SEM images of fabricated pillars before and after thinning with 5-min and 10-min BOE wet etch. C) SEM image of a nanopillar electrode. D) SEM image of a nanocrown electrode. E) SEM image of a Focused ion beam (FIB) milled cross section of nanopillar electrodes. F) FIB/SEM image of nanocrown electrodes (the Pt metal crown is outlined in orange and the SiO2 in grey). G) Schematic illustration of the interface of nanopillar and nanocrowns with the cell membrane. The membrane wraps tightly around nanocrown electrode arrays and interfaces with the SiO_2_ core. H) Impedance of nanopillar (blue) and nanocrown (green) NEAs at 1kHz. Nanocrown electrodes show lower impedance.I) Brightfield microscope image of a monolayer of iPSC-CM cells on a NEA device. J) Confocal images of a cell on a nanocrown electrode, focused on either the top or bottom of the nanocrown. The cell membrane (CAAX:green) and Integrin adhesion proteins (Anti-Itgb1:purple) can be seen on top of the nanocrowns (white arrows show location of nanocrowns). K) Higher magnified image, and a z projection of a cell on top of nanocrown electrodes showing the membrane (green) and integrins (purple) projection along the nanocrown height.

In the second part of the device fabrication, the Pt and SiO_2_/SiN_x_ -coated nanopillars were fabricated into either nanopillar electrodes or nanocrown electrodes (**Fig. 1A**). To fabricate nanopillar electrodes, we used a photoresist-masked dry etching step to selectively remove the insulation layer from the vertical nanoelectrodes. To fabricate nanocrown electrodes, the substrate was first coated with a thick and protective photoresist layer followed by a directional dry etching step to remove both the SiO_2_/SiN and the Pt layers from the tip of the nanopillars. Then, the substrate is subjected to a selective and time-controlled wet oxide etching to shorten the SiO_2_ core and to remove remaining insulating layers from the pillar. Scanning electron microscopy (SEM) images show the shape of nanopillar (**Fig. 1C**) and nanocrown (**Fig. 1D**) electrodes. In these images, the crown top of the nanocrown electrode can be clearly identified, with the conductive platinum layer appearing bright while the insulating SiO_2_ core appears dark.

To accurately examine the depth of the crown top, we used focus-ion-beam (FIB) to vertical mill open nanoelectrodes and then used SEM to image their vertical cross sections. From the FIB-SEM cross-section images, the nanopillar electrode shows a uniform platinum coating layer (**Fig. 1E**), while the nanocrown electrode has a hollow platinum crown 180-200 nm deep for the 3 μm-tall structure (**Fig. 1F**). We refer to these electrodes as “nanocrowns” to reflect on their shape and the irregular crown rim. Nanocrown electrodes have a lower impedance than nanopillar electrodes (**Fig. 1H, Suppl. Fig. S1**), likely due to the crown rim that is exposed to the electrolyte on both sides. Compared to the nanopillar nanoelectrodes, the nanocrown shape will induce the cell membrane to wrap around the outer surface while, at the same time, promotes cell adhesion to the inner core (**Figure 1G**). This new shape stabilizes the membrane-electrode interface and is important for the repeated and multiple-day recordings. hPSC-CMs were obtained according to our previous protocols (Burridge et al. 2014; Ye et al. 2021). hPSC-CMs cultured on the NEA devices exhibit spontaneous and rhythmic beating after 2-3 days and can be maintained on the devices for three months or longer (**Figure 1I**).

We used fluorescence imaging to examine cell adhesions to nanocrown electrodes. For this purpose, we transfected U2OS cells with a membrane marker GFP-CAAX and plated them on an NEA device (**Figure 1J**). Unless nanopillar is mentioned specifically, NEA refers to nanocrown electrode array in the subsequent studies. The cells were fixed and immunostained with anti-integrin beta1 to probe cell adhesions. A bright green dot around the nanocrown electrode indicates that the membrane is wrapping around the vertical nanoelectrodes. The magenta dot around the nanocrown electrode indicates the formation of integrin-containing adhesion complexes. It is worth noting that cell adhesions form on most but not all nanoelectrodes wrapped by the cell membrane. Confocal imaging in the z-direction shows the formation of cell adhesions at the top of the nanoelectrodes (**Figure 1K**).

As the new fabrication protocol involves photolithography and is wafer-based, the device throughput and reproducibility are significantly enhanced. Twelve NEA chips fit onto a single 4’’ wafer and more than a hundred NEA chips can be fabricated in a batch (**Suppl. Fig. S2)**. With each chip consisting of 58 NEA recording pads and two stimulation pads, these NEA devices increase the throughput by enabling parallel recordings and screening across many cell cultures and conditions as we show below. Since we used a SiO_2_ substrate which is effectively a transparent glass surface, these NEAs are compatible with optical imaging and long-term culture of hPSC-CMs.

### Nanocrown electrodes enable parallel, reliable and repeated iAP recordings

Electrophysiology measurements were carried out 5 to 50 days after cardiomyocytes were seeded on NEAs, when cells were in a confluent monolayer and beat synchronously. Before electroporation, NEAs reside outside the cell membrane and record eAPs with a biphasic spike shape and an average amplitude of about ∼1 mV, which is similar to that of planar microelectrodes. Upon the application of a single 200-µs electroporation pulse at ±3.5 V to a selected nanoelectrode, the recorded action potential transits from an extracellular spike shape to an intracellular waveform (**Fig. 2A**). When screening the electroporation pulse amplitude ranging from ±1 to ±5V, a ±3.5V electroporation pulse resulted in eAP to iAP transition with the highest amplitudes for nanopillar electrodes (**Fig. S3**). In the raw data trace in **Fig. 2A**, the electroporation event can be easily identified by a sudden rise and temporary saturation of the amplifier. In the same culture, three other nanoelectrodes were simultaneously selected for electroporation. Recordings from these channels showed intracellular recordings while their immediate neighboring channels showed extracellular spikes, indicating that the electroporation and the iAPs were highly localized to the selected nanoelectrodes (**Suppl. Fig. S4**).

**Figure 2.**
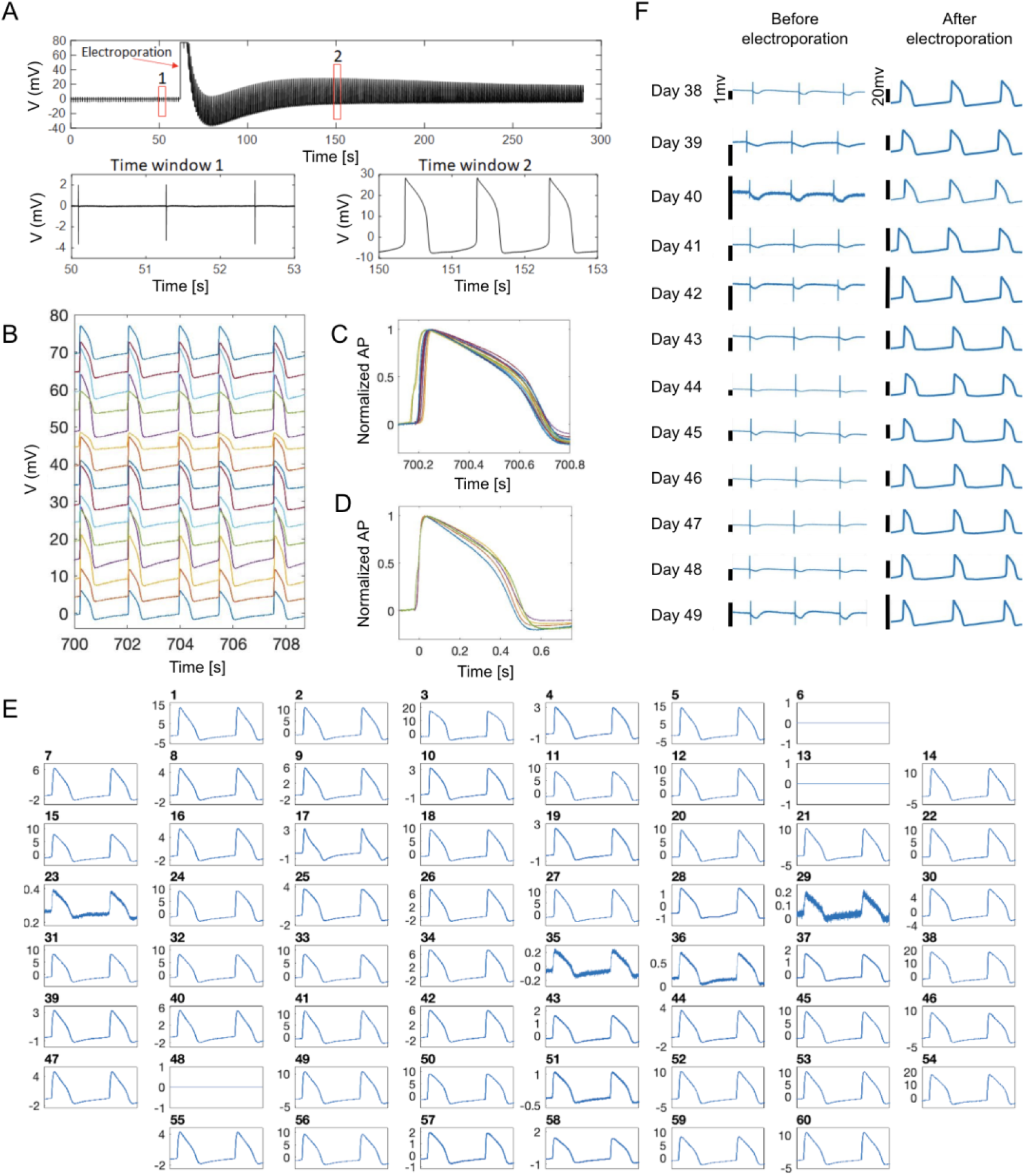
NEAs enable reliable and noninvasive recording of intracellular action potentials (iAPs). A) A representative raw trace of NEA recording from a hiPSC-CM. NEAs measure eAP spikes before electroporation (time window 1), and high amplitude iAPs waveforms after electroporation (time window 2). B) Simultaneous iAP recordings from 16 NEA channels show synchronous beating of hiPSC-CMs in the same culture. The 3-second traces are vertically shifted for clarity. C) Overlay of 16 amplitude-normalized iAPs illustrates 40-ms phase shift in depolarization time. D) Five of the 16 iAPs are aligned to their rising times. They show distinct iAP durations and waveforms for hiPSC-CM in the same culture. E) Simultaneous iAP recording from all NEA channels at t=15 min after many rounds of manual electroporation (excluding channels 6 which is defective, and 13 & 48 which are not NEAs). F) Twelve recording sessions of eAPs and iAPs from the same cell over two weeks in culture. The recording session started at day 38 on device. For each recording session, eAPs were first measured for 1 min before an electroporation pulse was applied, which converted the signal to iAPs. Each recording session lasted < 30 min and the culture was returned to the incubator afterward. It is clear that electroporation-induced membrane pores reseal after every measurement and eAP signals were detected for each recording session.

The electroporation-induced extracellular-to-intracellular transition rate is >99%, i.e. almost all eAP signals can be converted to iAPs after a single electroporation pulse delivered through the stimulator/amplifier system (Multichannel Systems, MEA1060-Inv). The amplifier system that we use has limited charge injection capacity and usually limits simultaneous electroporation to 4-6 cells. We will discuss a custom-built amplifier system that allows sequential electroporation from all electrodes in the last section of this article. Multiple rounds of manual electroporation applied to different sets of electrodes allow more iAPs to be recorded at the same time. In a set of experiments shown in **Fig. 2B**, 15 cells’ iAPs were recorded simultaneously. While their beating intervals were identical, these iAPs show differences in their amplitudes, upstroke times, and iAP durations. Amplitude-normalized traces show ∼49 ms phase shift in the iAP upstroke time (**Fig. 2C**). From traces recorded at spatially distinct electrodes, we calculated that the electric signal propagated in the culture dish with a velocity about 0.21 m/s (**Suppl. Fig. S5**). When these iAPs are aligned to their depolarization upstroke with normalized amplitudes, they show distinct waveforms and durations despite beating synchronously (for clarity, 4 of the 15 traces were displayed in **Fig. 2D**). The differences in waveforms are much larger when comparing iAPs from different cultures that are beating at different frequencies (11 iAP traces from 10 different cultures were shown in **Suppl. Fig. S6**). The differences in iAP duration, shape, upstroke slope, and repolarization form are indicative of inherent phenotypic variability among hPSC-CMs, which can be greater than drug-induced effects. Therefore, self-referencing of the same cell is important for assessing drug responses using hPSC-CMs.

In another set of experiments, we demonstrate that the NEA device allows simultaneous iAP recording of 57 hPSC-CMs. Before any electroporation (t=0-10s, Suppl. **Fig. S7**), all channels show extracellular signals except channel 6, which has a defective pin and is always grounded, and channels 13 and 48, which are connected to the two large pads designed for pacing as shown in Figure S2A. Then, we sequentially electroporated all 57 NEA channels that show eAP signals through multiple rounds of manual electroporation. At 15 min (t = 900-902s), all 57 NEA channels (100%) that were electroporated still show intracellular signals as shown in **Figure 2E**.

In addition to simultaneous iAP recordings, nanocrown electrodes allow daily and repeated recording of the same cell over weeks or longer. **Fig. 2F shows** high quality eAP and iAP signals over the time course of 12 days from the same cell. At each date, the culture was first measured for eAP recording, and then electroporated for iAP recording before being returned back to the incubator. In **Suppl. Fig. S8**, we show recordings from a cell that was repeatedly electroporated and measured for 15 times from day 3 to day 43 after being seeded on the device. These results indicate that long-term iAP monitoring is possible by repeated electroporation and recording sessions.

The extracellular-to-intracellular transition induced by electroporation is due to subnanometer-sized membrane pores that electrically connect NEAs to the intracellular domain. NEA-induced membrane pores are highly localized and reseal over time, during which the recorded signal can appear as a combination of intracellular and extracellular features (**Supp. Fig. S9**). We do not consider these juxtacellular signals as intracellular APs. Modeling of the electric fields shows that, during the electroporation step, the electric field is significantly enhanced at the rim of nanocrowns (**Suppl. Fig. S10**). To probe the size of the membrane pore, we monitored the cellular entrance of calcein, a method often used to probe membrane integrity. Calcein is a membrane impermeable fluorophore about 1 nm in size. We found that calcein was able to get into cells when added 1 min before electroporation, but not when added 1 min or longer after electroporation (**Suppl. Fig. S11**). This result suggests that the membrane pore has resealed to be less than 1 nm at 1 min after electroporation, when our NEA recording of iAPs usually starts. These membrane pores do not perturb the cell’s electrophysiology properties as we show later.

### The crown depth critically affects NEA performance

Our results indicate that the depth of the crown significantly affected the electrode performance. In our fabrication process, the depth was controlled by the SiO_2_ core etching time. We fabricated nanocrowns of two depths and systematically compared the performance of nanopillar electrodes, 450-nm-deep (∼10min etching time) nanocrown electrodes, or 180-nm-deep (∼3 min etching time) crowns. For this study, we recorded 76 cells using nanopillar electrodes, 111 cells using 450-nm deep nanocrown electrodes, and 308 cells using 180-nm-deep nanocrow electrodes, each for a 240s recording. These measurements were from many independent cultures and pooled together for statistical analysis. We define the success rate as the probability of recording intracellular signals shortly after a single +-3.5V electroporation pulse, on any cell that exhibits extracellular signals before electroporation (the recording and electroporation protocols and the device model are described in the method section). Both 450-nm-deep and 180-nm-deep nanocrown achieved >99% success rate, which was better than the 94% for solid nanopillar electrodes (**Fig. 3A**). The iAP amplitude measured by 180-nm-deep nanocrown electrodes (21.76 ± 9.95 mV, N=308) is twice as large as that measured by 450-nm-deep nanocrown electrodes (9.7 ± 8.13 mV, N=111) or by nanopillar electrodes (10.7 ± 9.22mV, N=76) (**Fig. 3B**).

**Figure 3.**
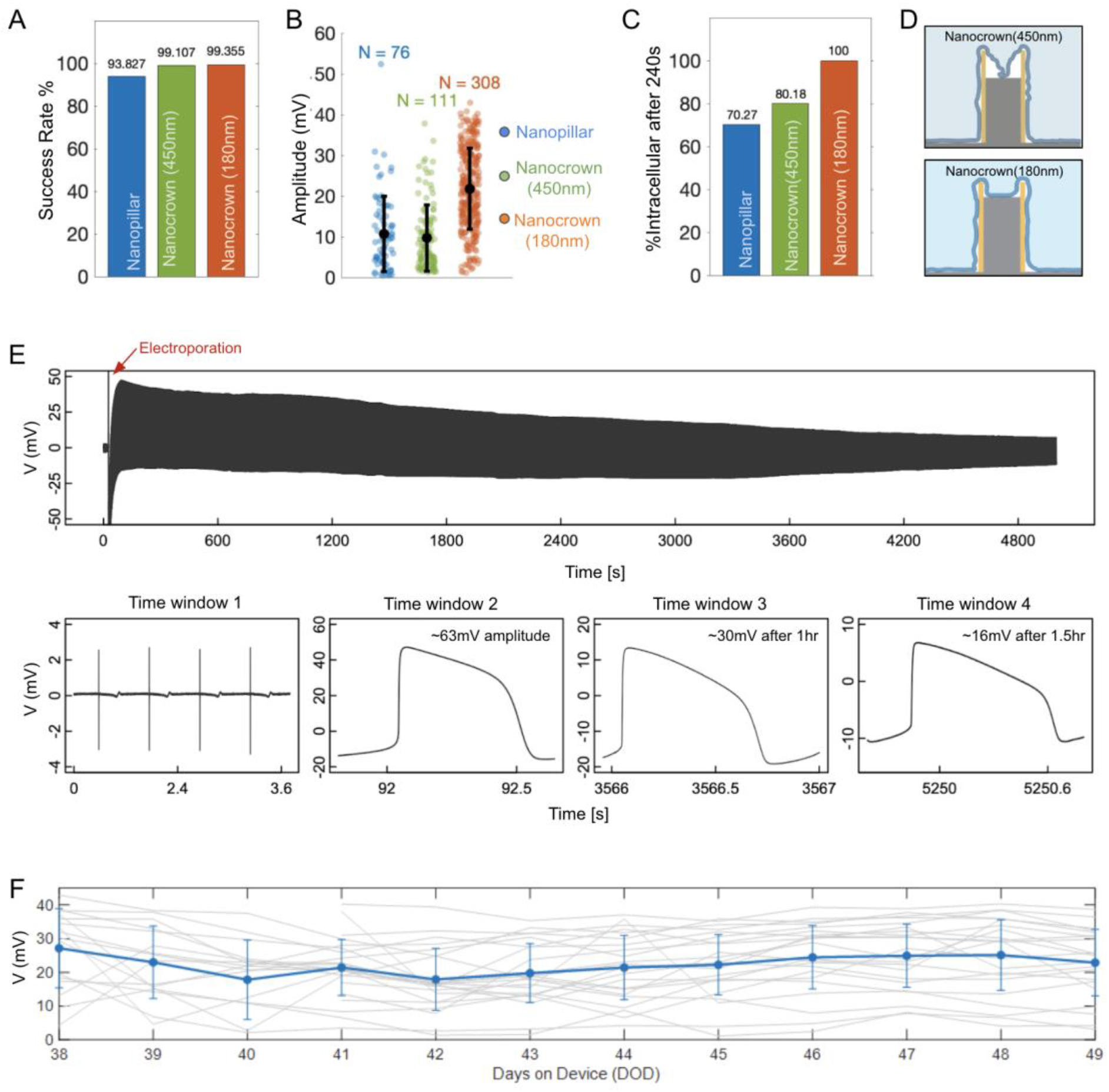
The crown depth critically affects the NEA performance. A) The initial amplitudes of iAPs measured by Nanopillar electrodes (blue), Nanocrown electrodes with 450nm depth (green) and Nanocrown electrodes with 180nm depth (Orange) B) Electroporation success rates for Nanopillar and Nanocrown electrodes. C) Percent of cells that maintained intracellular access at the end of the recording (∼240s after electroporation). D) schematic representation of the interface of the cell membrane and the nanocrown electrodes of various depths (450nm and 180nm). E) An iAP recording trace with 180-nm-deep nanocrown shows an iAP amplitude of ∼63 mV at t = 1min, ∼30mV at t=1 hr and ∼16mV at t=1.5hr. F) The average iAP amplitude of many cells that are recorded daily remains stable over the 12-day period, indicating that daily-repeated electroporation and resealing cycles do not degrade the recording quality.

When we compare the time-dependent decay of iAPs, 100% of the 180-nm-deep nanocrown electrodes maintained intracellular access at the end of 240s recording, as compared to 80% for 450-nm-deep nanocrown electrodes and 70% for nanopillar electrodes (**Fig. 3C**). In another set of experiments, longer 15-min recordings were performed using the 180-nm-deep nanocrown electrodes (N=41). An impressive 100% of these recordings maintained intracellular at the end of 15-min recordings, which was rare for nanopillar electrodes. This prolonged intracellular access is crucial for pharmacological experiments as we show in the next section. The superior performance of the 180-nm-deep nanocrowns is likely due to the cell membrane forming more stable adhesions to the inner SiO_2_ core than that for 450-nm-deep nanocrowns (**Fig. 3D**). Although we mostly limit our recordings to 15 min to avoid adverse effects to the cells as the culture is recorded outside the incubator, we observe from a few long-recording experiments that the intracellular access duration can be well over an hour for some recordings. For example, the trace in Fig. 3E shows an initial iAP amplitude ∼63 mV, which decreases to ∼30 mV at 1hr and is still >16 mV when the recording experiment was stopped at 1.5 hrs.

To systematically investigate how iAP signal changes when a cell is repeatedly electroporated and recorded over many days, we selected 16 cells from three cultures and measured the same cells everyday for a total of 12 days from day-on-device 38 (DOD38) to DOD49. Six more cells were added on DOD41 and recorded daily from DOD41 to DOD49. The maximum iAP amplitude is quantified for each recording trace and the amplitudes measured from the same cell are plotted as a gray line in **Figure 3F**. Our measurements show that the consecutive daily measurements do not decrease the average amplitude, which remains stable around 20 mV over 12 days (**Figure 3F, Figure S12**). These measurements confirm that daily-repeated electroporation and resealing cycles do not degrade the recording quality or perturb cells in the long term. Nanocrown electrodes are mechanically robust because their SiO_2_ cores are continuous with the bulk substrate. The nanocrown NEA devices can be repeatedly cleaned and re-used for many cycles of cardiomyocyte cultures. **Suppl. Fig. S13** shows the recordings from the same device in six independent cultures over a year. The device performance (signal amplitude and noise) is stable over repeated usage. This remarkable signal stability and device robustness afforded by nanocrown electrodes are not observed using other types of NEA devices.

### Simultaneous NEA and Patch recordings from the same cell confirm that NEAs record accurate iAP waveforms

To determine the accuracy of iAP waveforms recorded by NEAs, we compared these measurements with those recorded by manual patch clamp, which is the gold standard of intracellular recording. In these experiments, we performed simultaneous patch clamp and NEA recordings on the same cell (**Fig. 4A**). The transparent substrate allowed us to identify hPSC-CMs in contact with NEAs under a microscope for manual patch clamping (**Fig. 4B**). Upon obtaining a high seal resistance with the patch pipette, suction was applied through the pipette to breach the membrane and thus allowing iAPs to be recorded by the patch electrode. Before applying the electroporation pulse at t = 0, the patch electrode recorded typical iAP waveforms, while the simultaneous NEA electrode recorded characteristic eAP spikes (**Fig. 4C-D**). After NEA electroporation, both the patch and the NEA electrodes showed intracellular AP waveforms, albeit with different amplitudes (**Fig. 4D**). The electroporation event showed up as a sudden rise and saturation of amplifiers in both NEA and patch traces, which was used to align the two time traces. We note that sudden decrease of patch iAP amplitude due to loss of sealing resistance is common in patch recordings from cardiomyocytes that were not cultured on nanoelectrodes (**Suppl. Fig. S14**). Electroporation-induced decrease of patch iAPs is due to a reduction in the patch sealing resistance, not a change of the resting membrane potential, which would have dramatically affected the cell’s propensity to fire action potential and the refractory interval duration. As we show in the next section, the cycle time and the diastolic interval are not affected by the electroporation.

**Figure 4.**
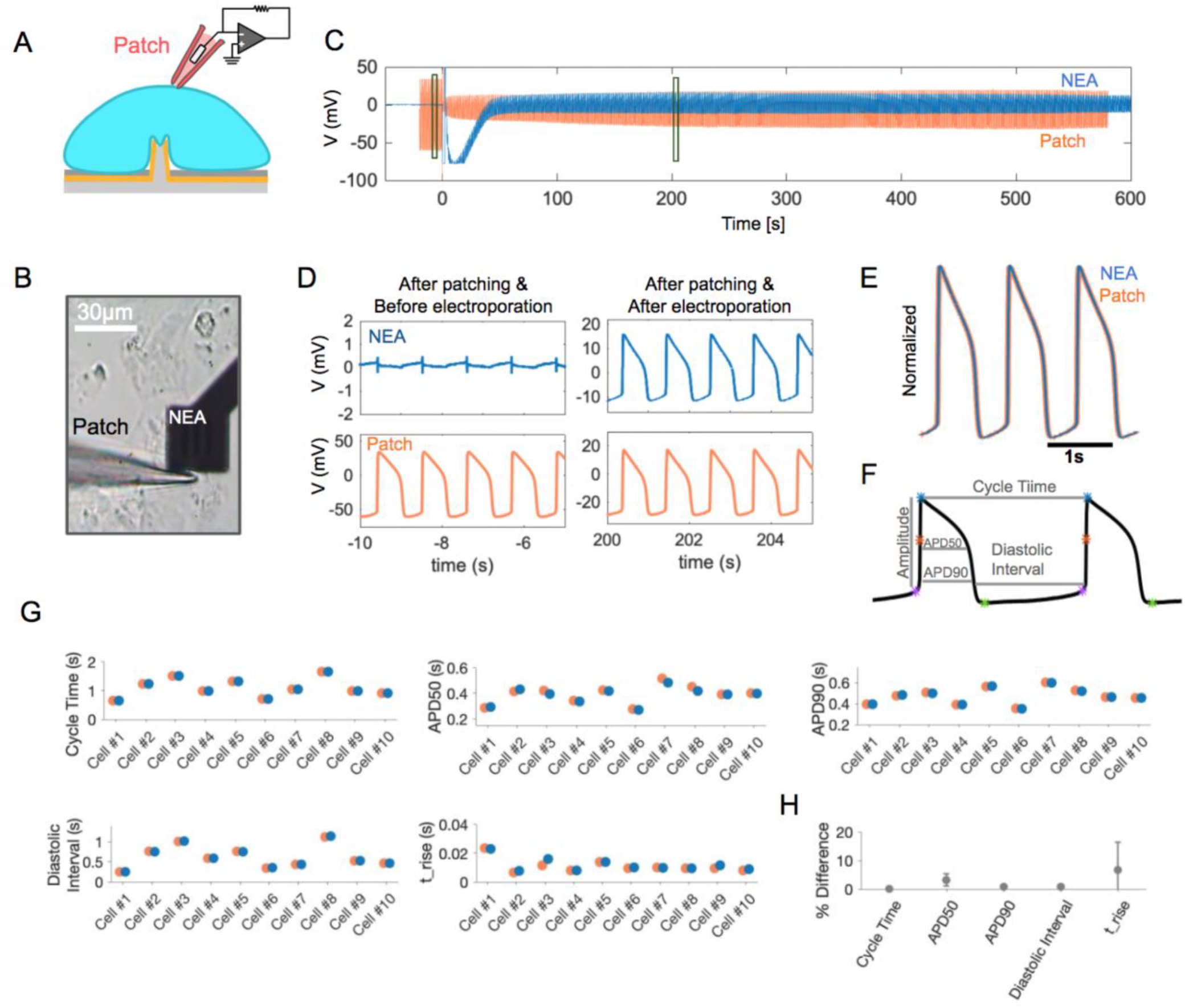
Simultaneous Patch clamp and NEA recordings verify the accuracy of NEA-recorded iAPs. A) A schematic representation of simultaneous patch and NEA recording from the same cell. B) A brightfield image showing a patch pipette approaching a cell on the nanopillar electrode from the left. C) Simultaneous recording traces by patch clamp (orange) and NEA (blue) from the same cell. The two traces are aligned in time with t=0 for electroporation. D) Zoomed-in time windows of NEA- and Patch-recordings before electroporation (left) and after electroporation (right). Before electroporation, NEA measured eAP spikes while patch clamp measured iAPs with high amplitudes. After electroporation, both NEA and patch measured iAPs. E) Three pairs of NEA- and Patch-recorded iAPs are amplitude normalized and overlaid together. The overlays show identical waveforms. F) Analyzed features of iAP waveforms: APD50, APD90, cycle time and diastolic interval and amplitudes. H) deviations of iAP features between NEA- and patch-recorded iAPs for ten independent cells. H) Averaged values of deviations of iAP features between NEAs and Patch for N=10 cells.

We wrote software to automatically calculate a single scaling factor for each pair of patch- and NEA-recorded iAPs to account for the difference in their amplitudes. The scaled and overlaid iAP traces show that the two waveforms almost entirely overlap (**Fig. 4E**). We carried out the simultaneous patch and NEA dual recordings for a total of 10 cells, all showing near-perfect overlap after scaling (see **Suppl. Fig. S15** for 8 more examples of dual-recording traces). From iAPs recorded by NEA and patch, we calculated a set of parameters known to be related to drug-induced proarrhythmia (**Fig. 4F**). These parameters included cycle time, diastolic interval (time interval where the membrane potential is more negative than 90% repolarization), action potential durations APD90 and APD50 (time taken to reach 90% and 50% of repolarization, respectively), and t_rise (the depolarization time from APD80 to APD20). Despite large variations among individual cells for each parameter, recordings of Patch and NEA from the same cell agree nearly perfectly with each other (**Fig. 4G**). Averaging over the 10 experiments, the differences between NEA and patch measurements were minimal: 0.05 ± 0.08 % for cycle time, 0.8 ± 0.6 % for APD90, and 3.21 ± 2.10 % for APD50, 0.85 ± 0.65 % for diastolic intervals, and 6.67 ± 9.75 % for t_rise (**Fig. 4H**). The comparison of simultaneous Patch and NEA recordings confirm that NEAs provide accurate measurement of iAP waveforms and parameters.

### NEA records accurate iAP waveforms despite time-dependent decrease of the iAP amplitude

Resealing of the electroporation-induced membrane pores over time causes the amplitude of NEA-measured iAPs to decrease over time. In the trace shown in **Fig. 5A**, the NEA iAPs decrease by an order of magnitude from 14 mV at 60 s to 1.3 mV at 530 s after electroporation. In the same time period, the patch iAPs from the same cell first increased and then exhibited large fluctuations after 430 s. The zoom-in plots show 15-s windows at three time points, t = 0, 80, and 430 s, for both patch and NEA traces (**Fig. 5B**). From the patch trace around t = 0s, we can clearly see that the amplitude of the patch iAP is reduced from 95 mV to 35 mV upon electroporation at t=0s. Despite the drastic change in the patch iAP amplitude, the cycle time is unperturbed and the overall iAP waveform remains the same. The NEA trace shows extracellular spikes before electroporation and then is out of range immediately after electroporation because of the amplifier saturation. In the t = 80s zoom-in window, the NEA electrode has recovered and both patch and NEA electrodes record iAPs that are synchronized in time and show similar waveforms but with different amplitudes. In the t = 430s zoom-in window, the patch iAP shows a sudden decrease in amplitude at t=432 s, likely due to a decrease of the patch sealing resistance. The event shows as a slight change of the baseline in the NEA channel without changing the iAP amplitude or the waveform.

**Figure 5.**
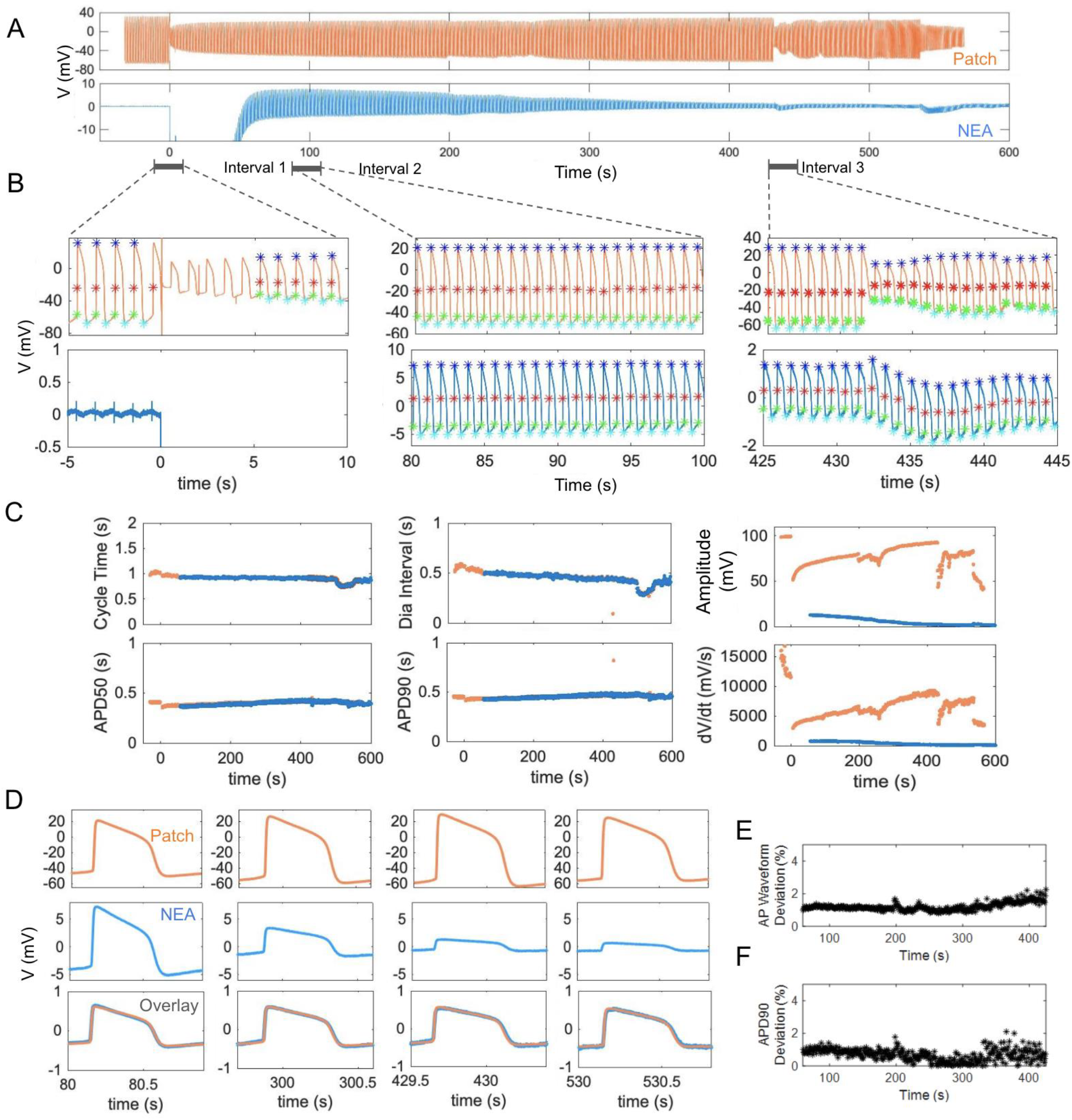
Comparison of APD features in simultaneous Patch clamp and NEA recordings. A) Simultaneous recording traces by patch clamp (orange) and NEA (blue) from the same cell. The two traces are aligned in time with t=0 for electroporation. D) Zoomed-in time windows of NEA- and Patch-recordings directly before and after electroporation (Interval 1), 80s after electroporation (Interval 2), and 425s after electroporation (Interval 3). The significant points on each iAP include spiking (red stars), starting (green stars), maximum (blue stars), and the minimum (cyan stars) points which are determined automatically by our in-house software. C) Overlay of time-dependant changes in APD features calculated using the Patch (Orange) or NEA (blue) recordings including cycle time (s), APD50 (s), APD90 (s), diastolic interval (s), amplitude (mV) and spiking velocity-dV/dt (mV/s). D) comparison of iAP amplitudes between patch (top-orange) and NEA (middle-blue) recordings at four time intervals after electroporation (80s, 300s, 430s and 530s). As the membrane pores sealed over time, the amplitude of NEA-recorded iAPs decreased. Nevertheless, the scaled NEA and Patch recordings (Bottom row) largely overlap at various time intervals. E) time dependent deviations of AP waveforms of patch and NEA recordings F) Time dependent deviations of APD90 calculated from patch or NEA recordings.

We developed a software package that automatically detects the spiking point (red stars), the starting point (green stars), the maximum point (blue stars), and the minimum point (cyan stars) of each iAP in the entire trace. A few iAPs immediately after the electroporation are omitted from analysis as there are often spurious artifacts. From these identified points, the software calculates the time-dependent evolution of cycle time, diastolic interval, APD90, APD50, amplitude, and spike velocity (dV/dt). Detailed analysis of traces shown in Figure 5A is presented in **Fig. 5C**. Detailed analysis of traces shown in **Figure 4C** is included in **Suppl. Fig. S16**. The analysis results show that the cycle time and the diastolic interval remain the same before and after electroporation despite the dramatic change of the amplitude. Therefore, the physiological functions of cardiomyocytes are not perturbed by NEA-delivered electroporation. APD90 and APD50 are slightly decreased immediately after electroporation but remain stable afterward. For the NEA trace, APD90 and APD50 remain constant for the entire time despite an order of magnitude decrease in amplitude. On the other hand, the amplitude and the spike velocity, which is dependent on the amplitude, show large changes. We note that, when the electroporation pulse is delivered simultaneously to many NEA pads, e.g. >30, it sometimes stimulates the whole culture and reduces the cycle time regardless of whether a cell is electroporated or not.

To further confirm that the patch and NEA iAP waveforms are not dependent on their amplitudes, we presented the scaled and overlaid iAPs at four different time points (**Figure. 5D**). During the time period, the patch iAP amplitude increased from 71.81 mV to 92.68 mV and then decreased to 81.39 mV. The NEA iAP amplitude decreased by about an order of magnitude from 12.76 mV (t=80s), 5.17 mV (t=300s), 2.28 mV (t=430 s), and 1.36 mV (t=530s). Despite their dramatic differences in amplitudes, the pairs of iAPs at the same time overlay nearly perfectly. The overall waveform deviation between paired Patch/NEA iAPs (**Fig. 5E**) and the APD90 deviation (**Figure. 5F**) are ∼1% for the entire period, which is negligible as compared to the drug effects we show later. From a simplified circuit model (**Suppl. Fig. S17**), both patch and NEA signals are linearly dependent on the biological signal, which explains why they report the same iAP waveform despite differences in amplitudes.

### NEA enables self-referencing assessment of drug-induced changes of iAPs

As shown in **Fig. S5** and in previous studies (Zhu et al. 2016), hPSC-CMs are mixed populations with cells showing heterogeneous iAP waveforms. Furthermore, the iAP waveform is affected by the cell density, beating frequency, and culture conditions (Du et al. 2015). The variability among iAPs underlied by phenotypical and physiological differences between cells and cultures can be much larger than drug-induced effects on a given cell’s iAP. This necessitates self-referencing - comparing the before-drug and after-drug iAPs in the same cell. Indeed, the standard patch clamp-based drug assay is self-referencing. However, the patch clamp method is labor intensive, low throughput, and not suitable for drug screening in hPSC-CMs.

NEAs enable self-referencing measurements of drug effects in parallel. For this study, we used NEAs to test a number of compounds with known arrhythmogenic risk selected from the CiPA library (Strauss et al. 2019; Colatsky et al. 2016) using common protocols in conventional pharmacology methods with patch clamp (Gibson et al. 2014). After electroporating and obtaining stable iAP signals, we first added DMSO as a vehicle compound (**Fig. 6A**). Subsequently, with continuous iAP recording, four increasing doses of a selected drug (dissolved in DMSO) were sequentially added to the cell culture every 3-4 minutes (0.3 nM, 1 nM, 3 nM, 10 nM of dofetilide for the trace shown in **Fig. 6A**). The mechanical agitation caused by liquid exchange induced signature fluctuations both in the measured channel and in the non-electroporated control channels, which we used to indicate the drug addition time. In each trace, 1000-2000 intracellular APs were recorded during the 15-20 min recording period (**Fig. 6A**).

**Figure 6.**
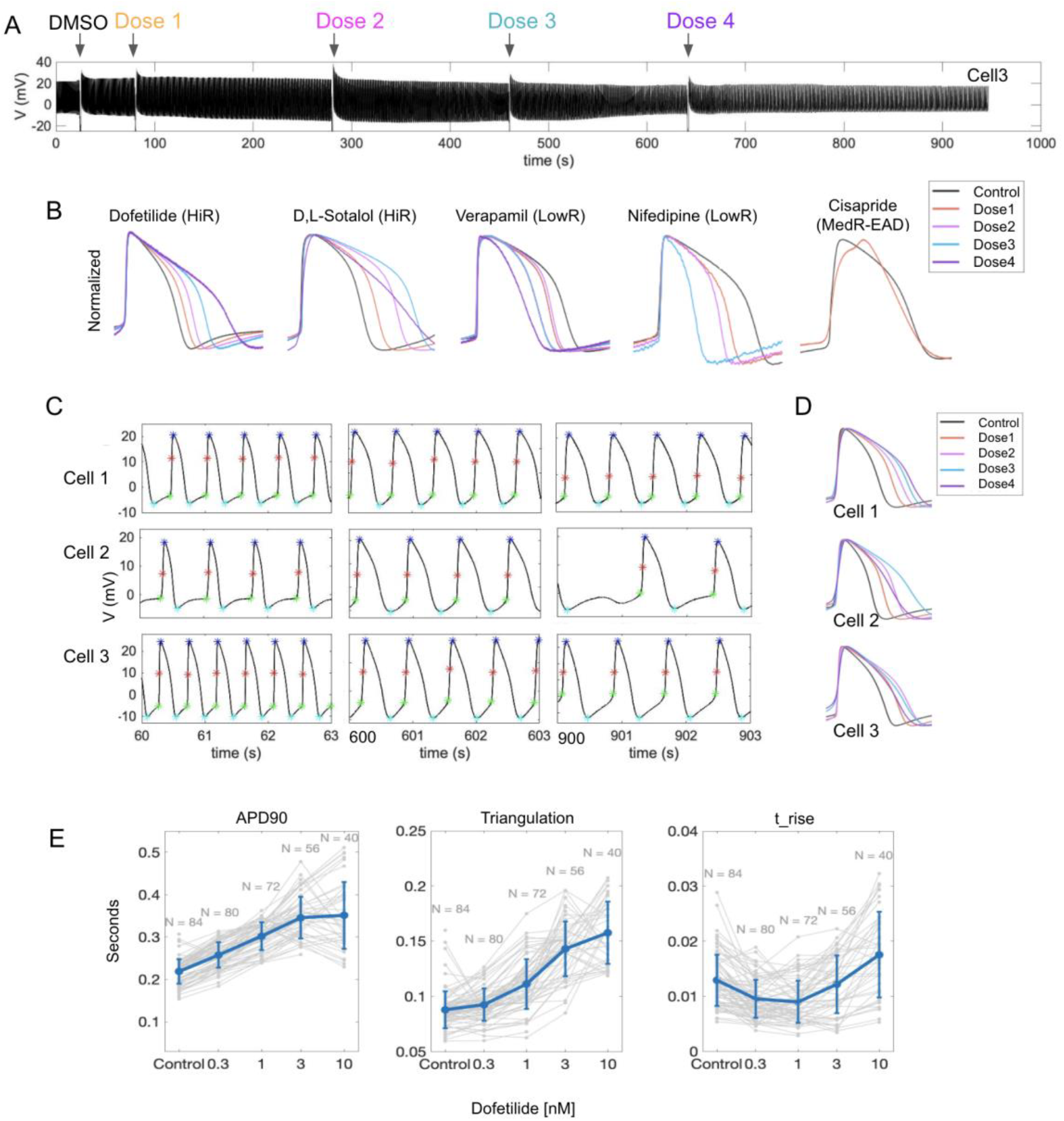
Self-referencing assessment of pharmacological compounds using nanocrown electrodes. A) A raw data trace of iAPs with sequential addition of pharmacological compounds (Dofetilide for the displayed trace). DMSO is added first as a control, after which an increasing dose of the selected drug is added every 3-4 minutes while iAPs are continuously being recorded. The large variations in the signal trace are due to vibrations associated with exchanging lipids when new doses of the drug are added (indicated by arrows). B) Overlay of amplitude-normalized and self-referencing iAPs, i.e. the same cell before and after adding increasing doses of each drug. The high risk drugs Dofetilide and D,L-Sotalol caused dose-dependent prolongation of iAPs, while the low-risk drugs Verapamil and Nifedipine caused dose-dependent shortening of iAPs. The medium-risk drug Cisapride caused early after-depolarization (EAD). C) Dose dependent changes in APD 90 for three cells showing distinct responses to Dofetilide. iAP waveforms at various time points during the pharmacology experiments at 60s (DMSO), 600s (Dose3), and 900s (Dose4) are shown. D) Self-referencing iAPs alignments for cells 1-3 show different dose-dependent responses to Dofetilide at doses 3 and 4. E) Self-referencing iAPs from 40-84 cells shows dose-dependent changes in APD90, APD90-30 (triangulation), and rising time in response to Dofetilide.

Using self-referencing NEA recordings, we compared a group of common high, low and medium risk compounds that are known to distinctly alter the shape of iAPs (**Fig. 6B**). High-risk compounds such as dofetilide and D,L-sotalol are known to prolong iAP durations, while low-risk compounds such as verapamil and nifedipine reduce iAP durations (Marschang et al. 1998; Bayer et al. 1977; Kimura et al. 1992). To examine drug effects, action potentials were selected for analysis 3-min after each dose addition. Action potentials from the same cell were normalized and aligned to their depolarization phase. By self-referencing, drug-induced prolongation of the action potential duration is clearly visible for dofetilide and D,L-sotalol (**Fig. 6B**). On the other hand, drug-induced shortening of the action potential duration is evident for verapamil and nifedipine (**Fig. 6B**). Another compound, cisapride induced early-afterdepolarization events. These data shows that NEAs are able to reliably capture drug-induced effects by self-referenced recordings.

Interestingly, after analyzing many cells, we found three types of dose responses to Dofetilide. This is represented in Figure **Fig. 6C** which shows 3 second zoom-in windows of the raw iAP traces at 60s, 600s, and 900s. For the first response type, increasing doses of dofetilide at 0.1, 0.3, 1, 3, 10 nM cause a monotonic dose-dependent increase of APD90 with the iAPs getting wider with increasing drug doses, but the fast uprising of the depolarization phase remains similar (represented ∼50% of cells).. For the second type that represented 30% of cells, the highest dose caused shortening of APD90, i.e. refractory dosage at 10 nM. For the third type, the medium dose of 3 nM already caused shortening of APD90, i.e. refractory dosage at 3 nM. For the second and the third types after the refractory dosage (10 nM for the second type and 3 nM for the third type), the iAP duration gets narrower but the shape is drastically different from the narrower iAPs at lower doses. In particular, the uprising duration is longer and the overall waveform is more triangularized. For cell 2, some action potentials are missing after 900 s.

Quantification of APD90, APD50, t_rise, and triangulation (APD90-APD30) highlights the variabilities of drug-induced responses in different cells and the importance of self-referencing in identifying these variations (**Fig. 6E**). The unique advantage of NEAs for drug assessment is that one can measure iAPs at multiple drug doses from the same cell and measure the responses of multiple cells in the same experiment. In this way, a set of dose-response data can be obtained with self-referencing to avoid the large cell-to-cell and culture-to-culture variations inherent to hPSC-CMs. APD90 and APD50 are often used for assessing drug effect. In a couple of days, we were able to measure 84 self-referencing dose-response curves using independent cultures (**Fig. 6E**). Membrane pores on some of these cells were sealed before higher doses were added, so there are fewer data points for higher doses, for example, N=84 for the control and N=40 for the highest dofetilide concentration at 10 nM (**Fig. 6E**). Nevertheless, the N=40 per dose sample size is already significantly larger than previous studies reporting iAP dose-response measurements with N<10. From the APD90 plot, it is obvious that some cells show monotonic dose-dependent increase, while others show refractory dosage at 3 nM or 10 nM. It is interesting that, for cells that show decreased APD90 after the refractory dosage, their t_rise parameter shows significant increase (**Fig. 6E**).

Unlike patch clamp or other invasive iAP recording methods in which the cells are no longer viable after recording, the NEA measurements are minimally invasive. Therefore, after washing out drugs and allowing 1-2 days of recovery in the incubator, the same hPSC-CM culture can be used again for assessment of the same or different drugs. In one experiment, we showed the dofetilide dose-responses from the same culture at day 1, day 3, and day 4, with the three independent sets of experiments showing slightly different drug effects induced by dofetilide (**Suppl. Fig. S18**). The fact that the same hPSC-CM culture can be used repeatedly for multiple drug assessment experiments reduces the number of precious hPSC-CM cell cultures required for obtaining dose-response curves, thereby facilitating the drug screening process.

### *In-situ* recording using miniaturized Petri-dish-like recording stage

All the data shown earlier were collected using a commercially available amplifier (MEA 1060, MCS), which requires that the culture be taken out of the incubator and placed on the recording stage for electrophysiology measurements. The health and the electrophysiological properties of cardiomyocytes are sensitive to experimental conditions such as temperature, humidity, and CO2 level (Kohlhardt and Haap 1976; Cedrini and Alloatti 1979). To reduce environmental variabilities, we recently collaborated with Cyion Technologies to design a miniaturized recording stage for in situ long term recording inside a CO2 incubator, which is controlled through a cable connected to a computer outside the incubator. The printed circuit board hosting our NEA devices was fitted with a bottomless 35mm Petri-dish **(Fig. 7A-B)**. In this design, the Petri-dish is placed on the recording stage with a short distance between the cell and the amplifier to minimize signal loss or distortion (**Fig. 7C**). In situ recording significantly reduced temperature-dependent variations in the cells’ electrophysiological behavior. Immediately after a culture was taken out of the incubator and mounted on a recording stage, we often observed a significant change in the cycle time, which is known to be affected by temperature (Cedrini and Alloatti 1979) (**Fig. 7D**). By comparison, the cycle time was much more stable when using our custom recording stage housed inside the incubator. The Cyion amplifier stage and custom-made software also supports automated sequential electroporation of all NEA channels in <1s with the potential for simultaneous iAP recording of up to 60 channels. Therefore, the miniaturized Petri-dish-like recording stage is ideally suited for iAP recordings. These new capabilities further enhance the advantage of using NEAs for preclinical evaluation and screening of new drug candidates in hPSC-CMs.”

**Figure 7.**
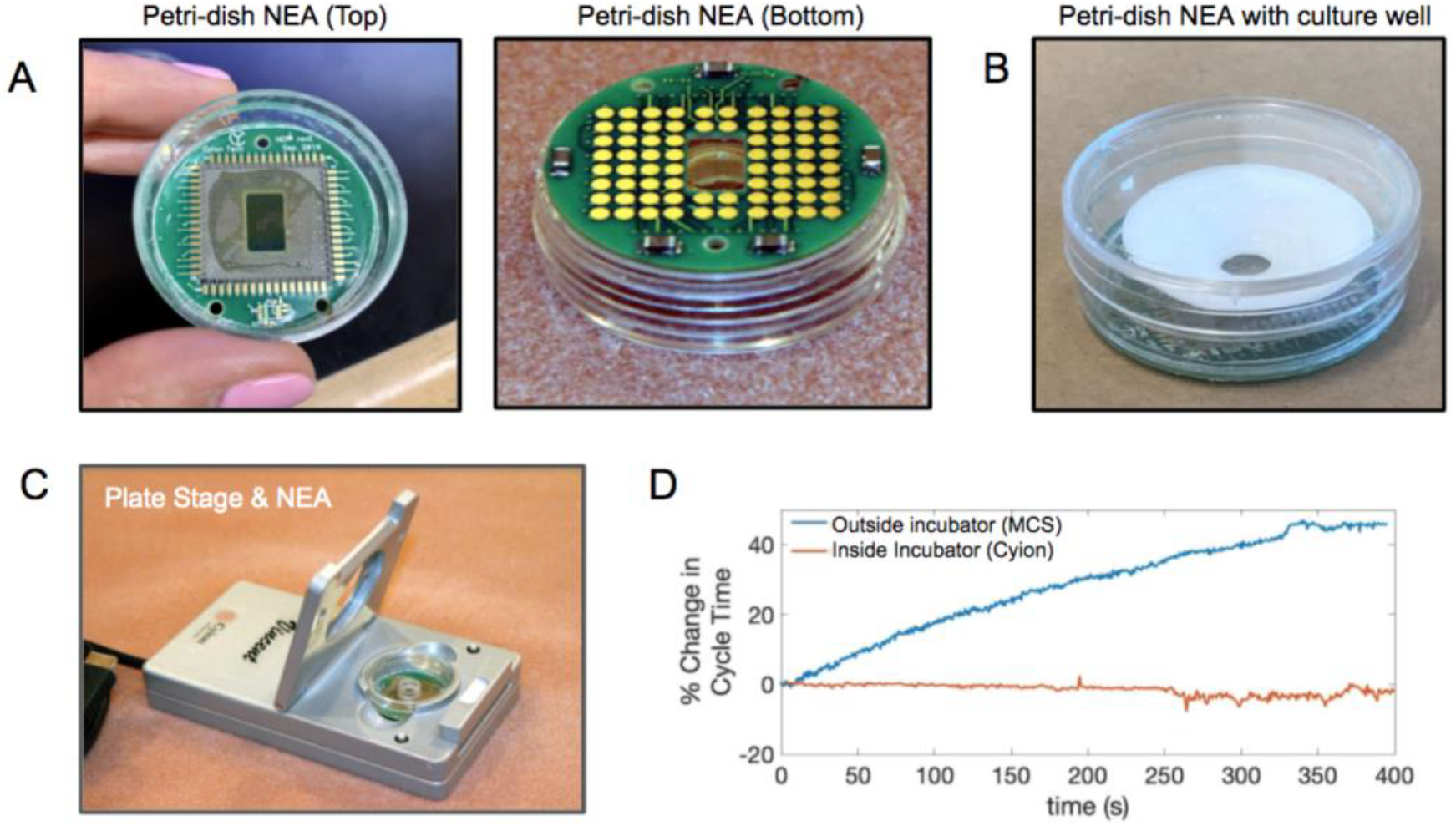
*In-situ* iAP recording using miniaturized petri-dish-like recording stage. **A)** Photographs of a Nanocrown device bound to a printed circuit board and integrated with a bottomless 35-mm petri dish. When placed on the recording stage, the electrode pads at the bottom are in contact to pogo pins inside the stage. **B)** A plastic 3D printed well (white) is assembled on top of the petri-dish NEA for cell culture. **C)** The petri dish NEA is situated in a Cyion recording stage. The recording stage is housed inside a CO_2_ incubator for *in-situ* recording. D) The measured cycle time of beating hiPSC-CMs gradually increases over 400s on the MCS recording stage (blue trace, outside the incubator), while the measured cycle time remains stable when using the miniaturized Cyion stage (orange trace, inside the incubator).

## Conclusions

Herein, we presented the development of semi-hollow nanocrown electrodes. The crown shape induces the cell membrane to wrap around the outer surface and to adhere to the inner core, which stabilizes the membrane-electrode interface. Nanocrown electrodes achieve 99% electroporation success rates and reliably measure iAPs from cardiomyocytes with high accuracy and signal strength. The prolonged recordings allowed us to perform a series of pharmacology experiments based on the self-referencing iAP waveform. The development of a Petri-dish NEA and an accompanying miniaturized recording apparatus would enable in situ recording of iAPs inside CO_2_ incubators. These developments will hopefully enable iAP-based assays for dose-dependent drug screening.

## Methods

### Nanopillar and Nanocrown electrode array fabrication

Quartz wafers were cleaned and deposited with HMDS to promote resist adhesion. A positive photoresist (Shipley 3612) was spin-coated and exposed by a maskless aligner (Heidelberg MLA-150) to pattern circular holes. A Cr mask of 120 nm was later deposited by metal evaporation before being lifted-off in acetone to make circular Cr disks. Using the deposited Cr as a mask, vertical microstructures were subsequently generated by anisotropic reactive ion etching with a mixture of C4F8, CHF3, and Ar (Versaline LL-ICP Oxide Etcher) to precisely control the vertical profiles of the quartz nanostructures. The substrates were then immersed in Chromium Etchant 1020 (Transene) to remove the chrome mask and subsequently wet-etched in Buffered Oxide Etchant (BOE) 20:1 (Transene) to decrease the nanostructures’ dimensions. For electrode fabrication, a bilayer resist strategy was used to construct metal connection lines with smooth edges. Microposited LOL and Shipley 3612 were spin-coated, baked, and then exposed to create an undercut in the Microposited LOL layer underneath the resist for the lift-off step. A layer of Ti (10 nm) and Pt (40 nm) was sputtered and then lifted-off in Microposit Remover 1165 for 16 hours. Subsequently, the insulation layers composed of 60 nm of Si3N4 and 540 nm of SiO2 were deposited via plasma-enhanced chemical vapor deposition (350 °C; Plasma-Therm Shuttlelock).

For nanopillar electrodes, the insulation layer was removed from the nanopillars through the spin coating of a thin layer of resist, followed by a wet etch in Buffered Oxide Etchant 20:1 (Transene). For nanocrown electrodes, a hollow conversion strategy was applied after the insulation step to create nanocrowns. Briefly, a layer of photoresist was spin-coated to protect the entire substrate. Then, the substrates were anisotropically etched with Ar (MRC model 55 RIE) to remove the insulation layer and the metal layer at the tip of the nanostructures and expose the inner quartz. The inner quartz core was subsequently etched with BOE The depth of the nanocrown was controlled by varying the etching time between 3 to 10 minutes. Nanocrowns with 450 nm or 180 nm crown depths were fabricated by 10 min or 3 min BOE etching respectively. Finally, the quartz wafers were diced into 12.5 mm x 12.5 mm pieces by the Micro Dicing Services to fit our custom made printed circuit boards. The final devices were characterized by SEM (FEI Nova).

### hPSC-CM differentiation

hPSC-CMs were differentiated and purified based on our previous publications (Wu et al. 2019; Burridge et al. 2014; M.-T. Zhao et al. 2017). Briefly, hPSCs (line SCVI-273) with 90-95% confluency were treated with 6 µM CHIR99021 (Selleck Chemical) in RPMI (Gibco®, Life Technology) supplemented with B27 w/o insulin for 2 days, and allowed to recover for 1 day in RPMI plus B27 without medium. Then cells were treated with 5 µM IWR-1 (Selleck Chemical) for 2 days. After recovering in the fresh RPMI plus B27 without insulin medium for another 2 days, cells were switched to RPMI plus B27 with insulin for 2 days. Beating hPSC-CMs were derived around 9-11 days after differentiation. The hPSC-CM were purified with glucose-free RPMI plus B27 with an insulin medium for 2-4 days. Differentiated hPSC-CMs were maintained in RPMI plus B27 with insulin medium for the following experiments.

### Immunostaining of cell adhesions

The samples were incubated with 1:100 diluted anti-activated integrin β1 antibody (Sigma-Aldrich, MAB2079Z) in HEPES-Buffered Hanks Balanced Salt Solution (HHBSS), for 30 min at 4°C. The samples were then washed with ice-cold HHBSS for 3 times and fixed with 4% Paraformaldehyde in PBS for 15 min at room temperature (RT). The fixed samples were washed with PBS for 15 min and incubated in blocking buffer (5% Bovine serum albumin in PBS) for 1 h at RT. The sample were then incubated with 1:500 Alexa Fluor 488-conjugated goat anti-Mouse IgG (H+L) secondary antibody (Thermo Fisher Scientific, A11001) in blocking buffer at RT for 45 min, washed with PBS for 15min at RT.

### Cell Culture of hPSC-derived-CMs on NEA devices

hPSC-CMs were maintained in Matrigel pre-coated 6-well plates in RPMI 1640 medium (Gibco REF 11875-093 500ml) with B-27 supplement (Gibco REF 17504-044 10ml). The culture medium was refreshed every 2-3 days. Before seeding cells on NEA devices, the devices were coated with 1mg/ml poly-L-Lysine at room temperature for 15min, 0.5% Glutaraldehyde in PBS at room temperature for 10min, and followed by 1:200 Matrigel (Corning REF 356231) in DMEM/F12 (Gibco REF 10565-018) at 37°C for 3 h or overnight. A three times PBS wash was performed between each coating step. Cells were dissociated with TryPLE select 10X (Gibco REF A12177-01) at 37°C for 5 min and ∼12 ×10^5^ cells were plated in each coated device in culture medium supplemented with 10% KnockOut Serum Replacement (KSR)(Gibco REF A31815-01), and cultured in a 5% CO2 incubator at 37°C. The medium was refreshed with normal culture medium every 1-2 days from the second day after replating. Cells are maintained in regular cell culture medium before and after electroporation and during recording. Measurements were done 5-30 days post-plating unless otherwise stated. The devices are reusable after the following washing steps in the device wells: 5 min of incubation with TrypLE Express (Gibco REF 12605-028) at 37°C to remove cells, overnight incubation with Enzymatic cleaner (Ultrazyme) in PBS at room temperature, and leave in 70% ethanol for at least 1 h. The wells are always immersed in 70% Ethanol until the next usage. The devices were dried of ethanol and further decontaminated with UV illumination for 30 min before coating for the next cell culture.

### Electrophysiology Recordings

All recordings were performed by a 60-channel voltage amplifier (MEA1060-Inv-BC, Multi-Channel Systems, Reutlingen, Germany), equipped with a plate heating system which was maintained at 37°C for all recordings. Recordings were performed at a sampling rate of 5 kHz. Simultaneous patch-clamp and NEA recordings were carried out on hPSC-CM at room temperature in the culture medium (RPMI 1640 + B-27). For patch-clamp, the action potential was recorded under whole-cell current-clamp mode using Multiclamp 700B amplifier and Digidata 1440A digitizer (Axon Instrument) with pClamp10 software (Molecular Device). Borosilicate patch pipettes (Sutter Instruments, O.D.: 1.5mm, I.D.: 0.86mm) were pulled with a P1000 pipette puller (Sutter Instrument) with resistances between 2 and 6 MΩ. Pipette internal solution contains in mM: 120 K Aspartate, 20 KCl, 1MgCl_2_, 5 EGTA-KOH, 0.1 CaCl_2_, 10 HEPES, 4 Mg_2_ATP. The pH was adjusted to 7.20 with KOH. The patch-clamp recording was done at 10kHz sampling rate and a 10kHz Bessel filter was applied. During the experiments, cells occupying NEA pads were located through a microscope. The extracellular signal recording was initiated on NEA followed by patching of the cell. After action potential signals were observed through patch-clamp, electroporation was applied to the same cell through nanoelectrodes to get the intracellular signal. Recordings from both NEA and patch electrodes started before the electroporation and continued during and after the electroporation. Data analysis was performed with Matlab and Python as stated below.

### Electric field models

To assess the difference in electric field strength between hollow and full nanopillars, we performed a finite-element analysis of the voltage distribution in 3D in vacuum. We did not simulate the electrolyte and double-layer effects for simplicity. Therefore, the obtained results can qualitatively translate only to the electric field strength during fast transients when operating the nanopillars in electrolytes. We used FEniCS, a Python implementation of the finite element method which allows simulating partial differential equations in complex geometries.

Exploiting the radial symmetry of the problem, the simulation reduces to 2D by simulating on a plane through the pillar’s center. The full nanopillar is simulated as a trapezoid to enable simulating angled wall profiles. The hollow nanopillar consists of the same trapezoid where the outer edges are thickened to a given value and the top edge is removed.

The weak form of Laplace’s equation in vacuum with no source term is solved on the domain which consists of a square that is 10 times the height of the nanopillar in every dimension in order to minimize numerical edge effects. The boundary conditions are 0 Volt on the box edge, except for the bottom of the nanopillar. Every edge of the nanopillar is set to 1 Volt.

The electric field is computed by taking the gradient of the resulting voltage field in the x and y directions. The magnitude of the electric field is obtained by summing the square of the x and y components of the electric field. By interpolating the magnitude of the field spatially, we can compare the relative electric field magnitudes between the full and hollow nanopillars at the same locations (**Fig. S9**). The code to generate the voltage and electric field distributions based on the FEniCS Python package is available at [DATA_AVAILABILITY_REPO] in the form of a Jupyter Notebook.

### Analysis of iAP data

All data analysis was automated and free of manual intervention. From a raw iAP trace recorded by either NEA or patch electrode, its corresponding standard deviation trace is calculated using a 1-ms sliding window. A peak finding algorithm subsequently identifies the peaks from the standard deviation trace, which correspond to the sharpest-rising points of individual iAPs. Then, the software examines the raw data trace to the left of each rising point to identify the starting point of each iAP, and to the right of each rising point to identify the maximum point of each iAP. The amplitude is calculated from the starting point to the maximum point. APD90 and APD50 are calculated as the width of the iAPs at 90 and 50 percent of repolarization. T_rise is the time that it takes from the 20 percent to the 80 percent of depolarization.

For scaling Patch and NEA recorded iAPs, the two recording traces are first time-aligned to the electroporation time and resampled to the same frequency of 2 kHz. Then, each trace is separately processed to identify the rising points of all iAPs. The software then automatically calculates a scaling factor for each pair of iAPs to account for their differences in amplitudes. This is done using the mean square error algorithm by first setting the mean of iAPs to 0 and normalizing the amplitude of the patch iAP to 1. Then, the scaling factor for the NEA iAP is calculated as sum(iAP2.*iAP1)/sum(AP2^1). After scaling, the deviation of the two iAP waveforms is calculated as the mean absolute deviation.

### Pharmacology experiments

All pharmacology experiments were conducted using the following procedure. First, 4-6 cells were electroporated and iAP signals were obtained and recorded until a stable baseline was reached for all signals. In order to obtain consistent recording files for all pharmacology experiments (such as the one shown in **Fig. 6A**), after the stable iAP baseline was reached, a new recording file was then started and continuously recorded for 30 min. At 20 s, DMSO was added as a control, followed by a sequential addition of increasing doses of the selected drug at 80 s, 280 s, 460 s and 640 s (as shown in **Fig. 6A**).

*Data analysis for Pharmacology experiments*. Individual peaks were detected at and around time 75s for the control data, and at 260 s, 440 s, 625 s, and 815 s for the four drug doses. The three action potentials closest to these timepoints were used for analysis. Differences in APD values between these three action potentials were usually under 1%. If AP signals at a low drug dose started to exhibit extracellular signatures, the data will not be analyzed for higher drug doses. To overlay the iAPs at different drug doses, we time-aligned the traces to their sharp rising point with normalized amplitudes.

### Data availability

The data that support the findings of this study are available from the corresponding authors upon reasonable request.

### Code availability

Matlab script for plotting out the raw data and time alignment is available from the corresponding author upon reasonable request.

## List of abbreviations

NEAs: Nano-pillar Electrode Arrays
AP: Action Potential
iAP: Intracellular Action Potential
eAP: Extracellular Action Potential

## Acknowledgements

This work is supported by National Institutes of Health (R01GM125737 (Cui), 1R35GM141598 (Cui), R01HL155282 (MTZ), R01HL145676 (JCW), and UH3TR002588 (JCW)), American Heart Association (AHA) Career Development Award 18CDA34110293 (MTZ), and Google LLC. award (ZJ and Cui).

