## Supplemental Figures for "Nanocrown electrodes for reliable and robust intracellular recording of cardiomyocytes and cardiotoxicity screening"

List of abbreviations

| | |
|---|---|
| NEAs | Nano-pillar Electrode Arrays |
| AP | Action Potential |
| iAP | Intracellular Action Potential |
| eAP | Extracellular Action Potential |

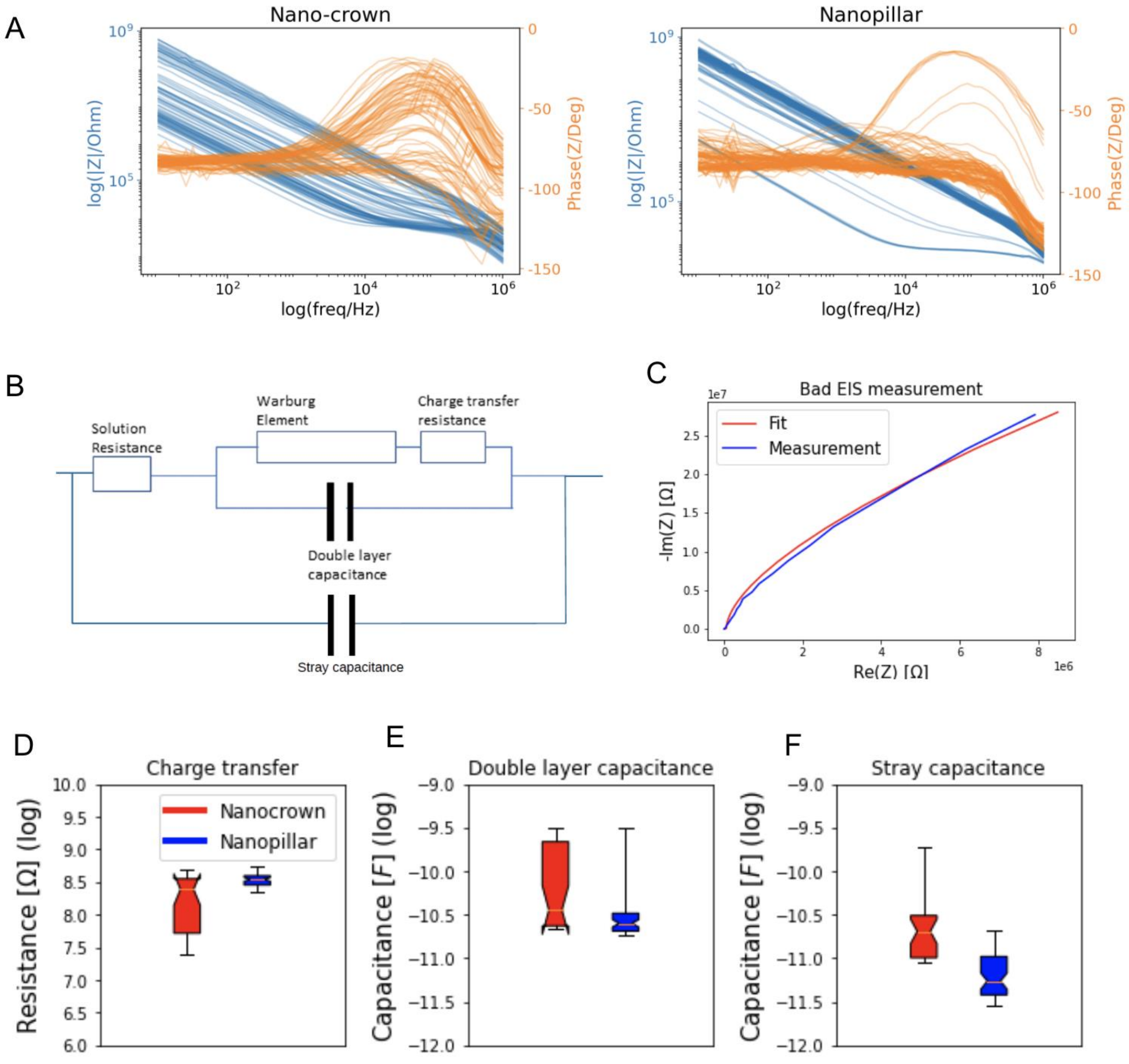


**FIgure S1** A) Impedance spectrum from 0.1Hz to 10MHz for nanocrown and nanopillar electrodes. B) Schematic of circuit used to model the EIS curve: Randles circuit in series with a capacitance. Randles circuit consists of a series of Warburg element and charge-transfer resistance, with both of the latter in parallel with a double layer capacitance. In addition to that, a resistance is connected to it in series to model the solution resistance. The addition of a capacitance in parallel with the Randles circuit models the stray capacitance. C) representative EIS measurement and modeled fit. Box plots of the fitted parameters along with their distribution (20%-80% percentile for **D)** Charge transfer, E) Double layer capacitance and F) stray capacitance as calculated by our model. N=85 EIS for nanocrown and N=100 for nanopillar.

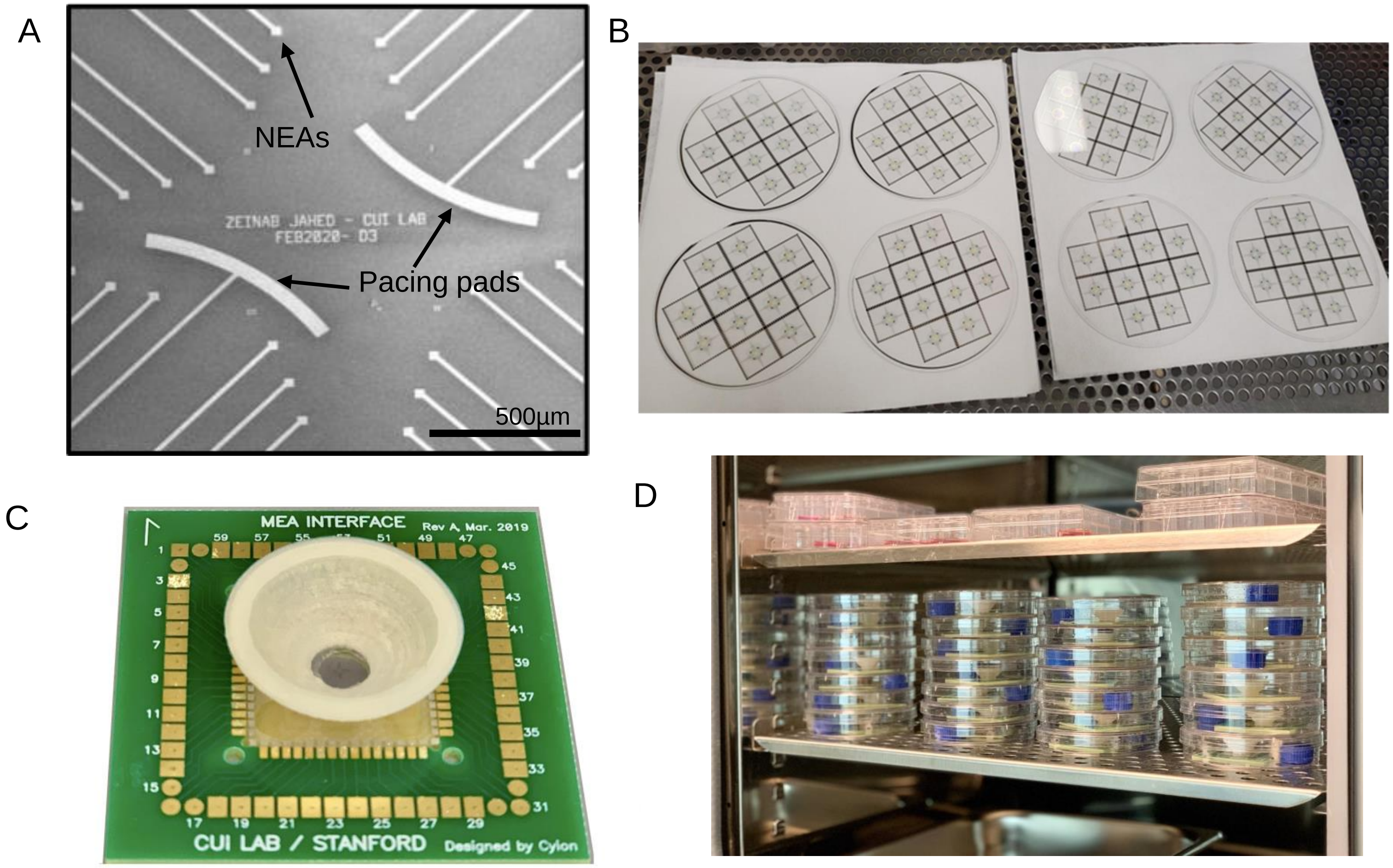


**Figure S2. NEA devices for cell culture.** A) design of NEAs including two large pacing pads B) NEA chips are wafer-based and batch fabricated in the cleanroom. C) NEA chip is bonded to a printed-circuit board for electrical measurements using an amplifier system by Multi-Channel Systems. The conical-shaped and 3D-printed cell culture well is glued to the center of the NEA chip. D) Assembled NEA devices were housed in 10-cm petri dishes inside a $CO_2$ incubator. A blue water container was included in each petri dish to maintain local humidity.

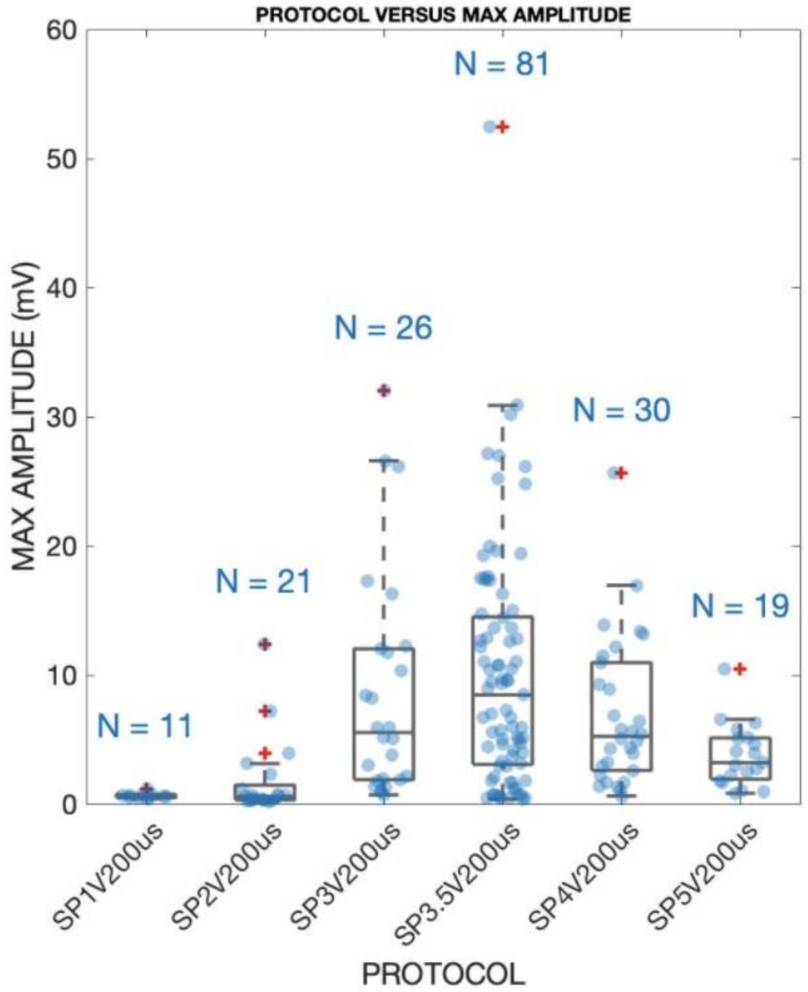


**Figure S3.** The effect of electroporation pulse amplitude on the maximum AP amplitude obtained with nanopillar electrodes.

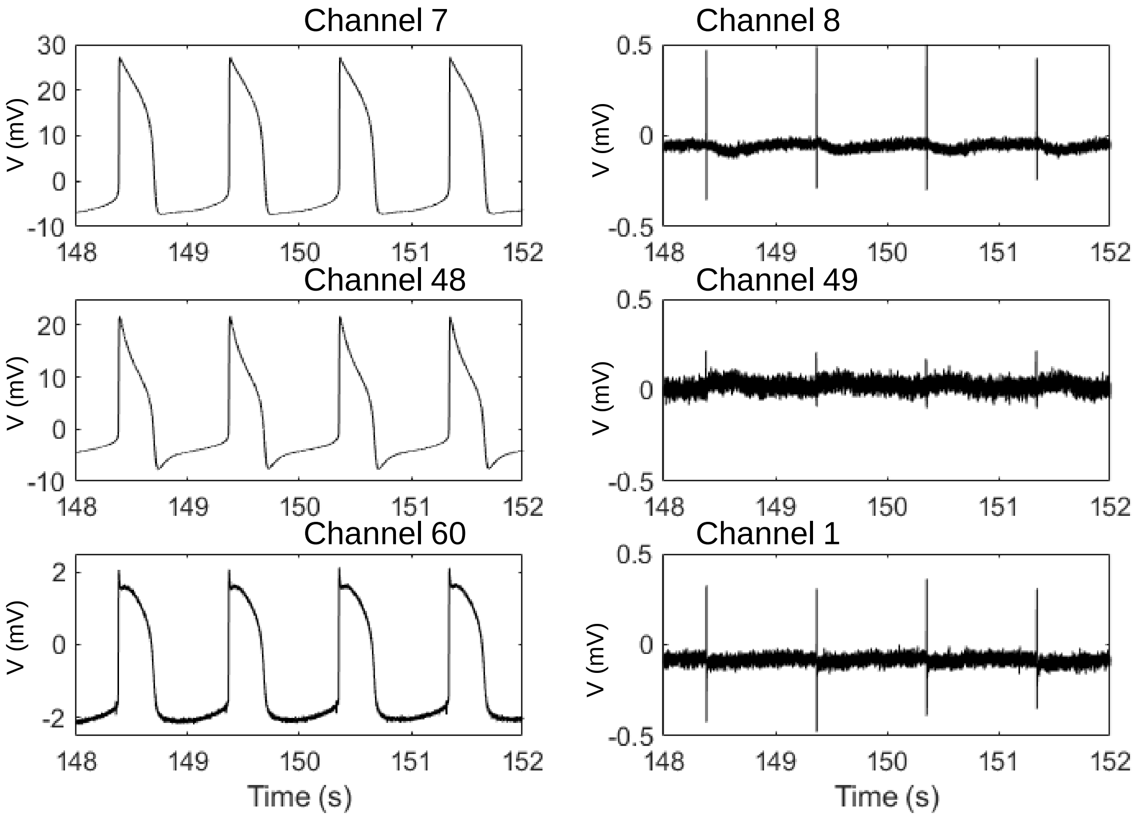


**Figure S4. Simultaneous recording of iAP and eAP signals from neighboring channels.**
Three pairs of neighboring channels in the same culture. The left channels were electroporated and recorded intracellular signals, while their neighboring right channels continued to record extracellular spikes at the same time range.

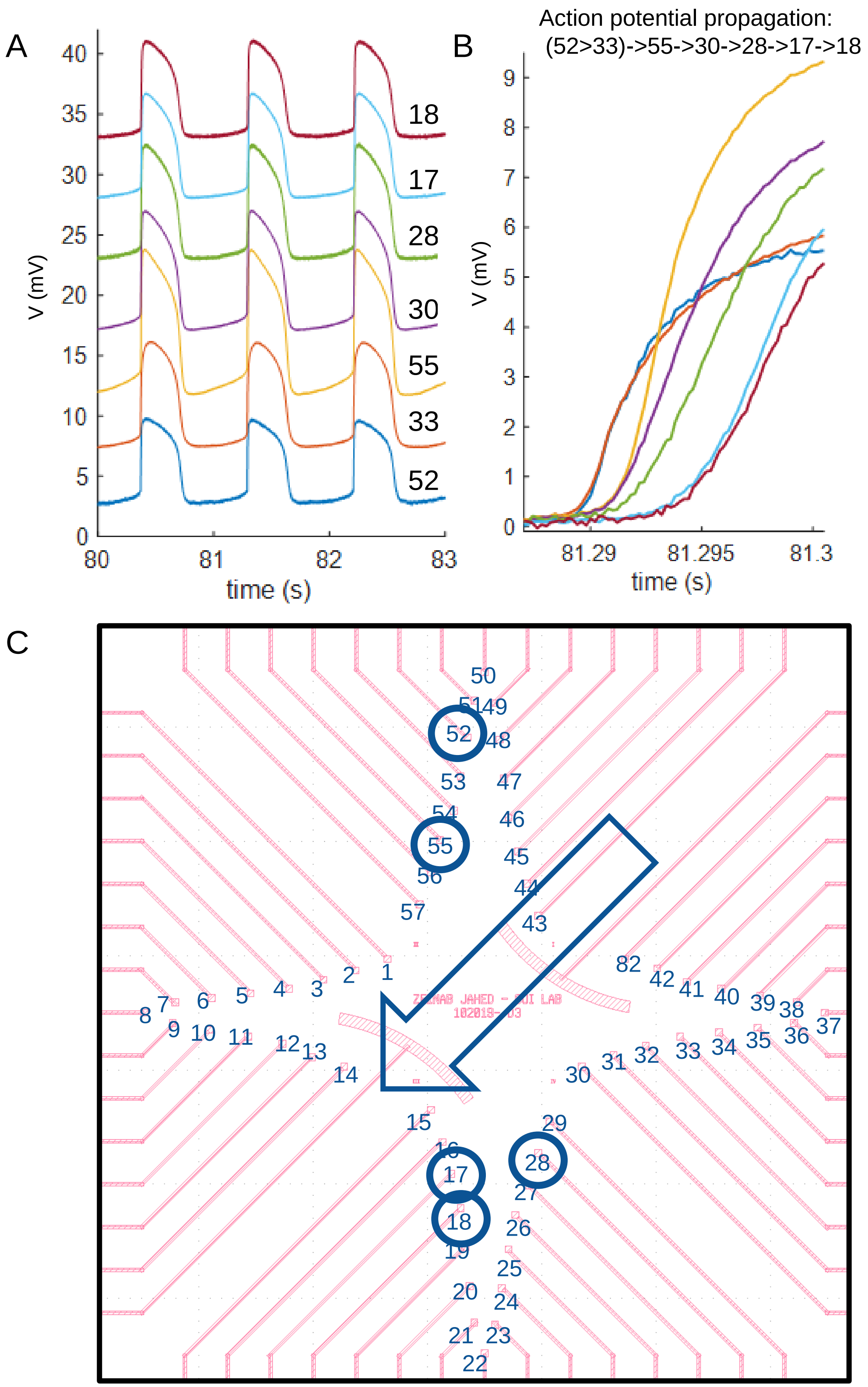


**Figure S5. Propagation of APs on NEAs.** A) simultaneous recording of iAPs from multiple channels within the same culture (The traces are vertically shifted and the amplitudes of three traces 18, 52, 28 are multiplied by 2 for visual clarify). B) Overlay of iAPs to show sequential depolarization of iAPs for cells in the same culture. C) Direction of AP propagation determined by measured depolarization times and the physical location of each channel on the NEA chip.

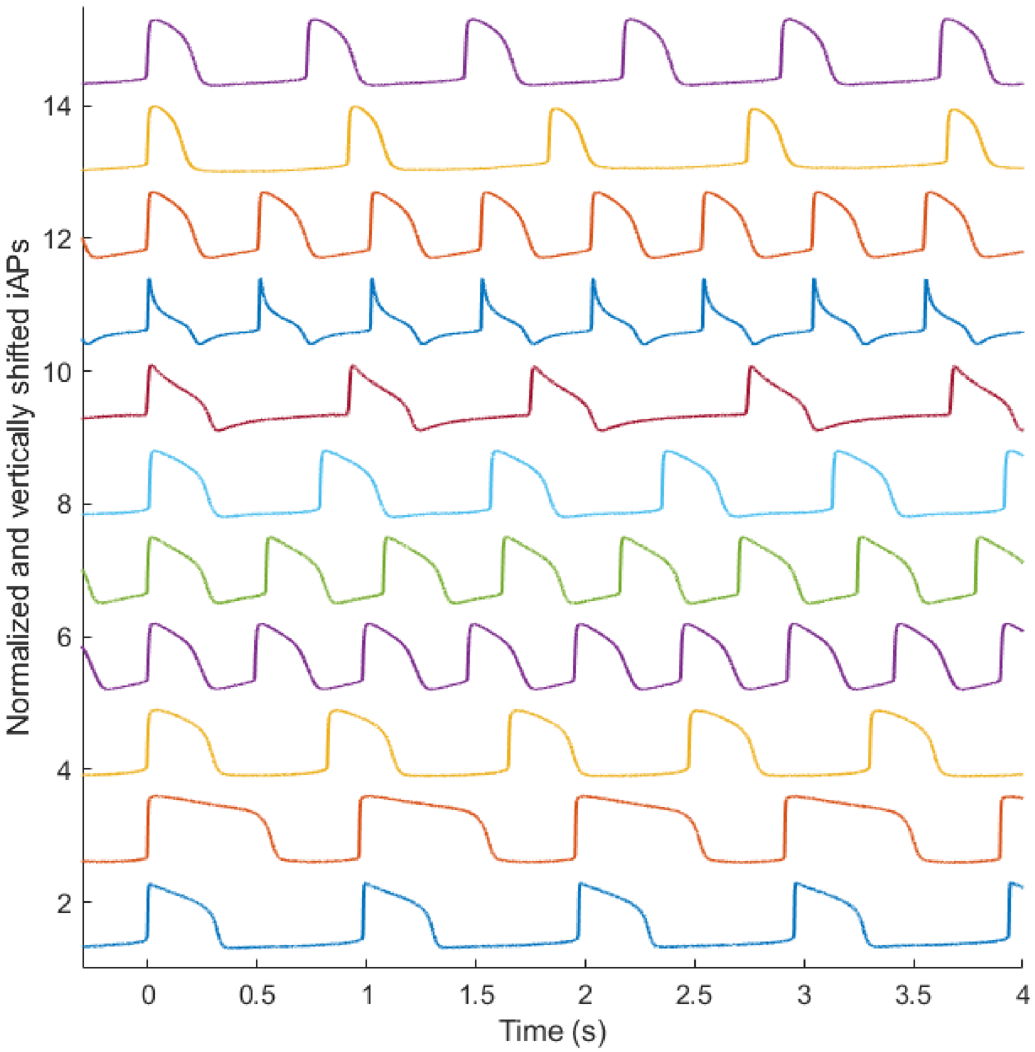


**Figure S6. Diversity of iAP waveforms.** 4-second recordings are shown for 11 iAP traces recorded from 10 different experiments. The iAPs are aligned in time to the rising phase of the first peak to visualize the changes in frequency and iAP waveforms. The amplitudes are normalized and the traces are vertically shifted for clarity. The lowest two traces are from the same culture, showing identical beating intervals but very different iAP durations.

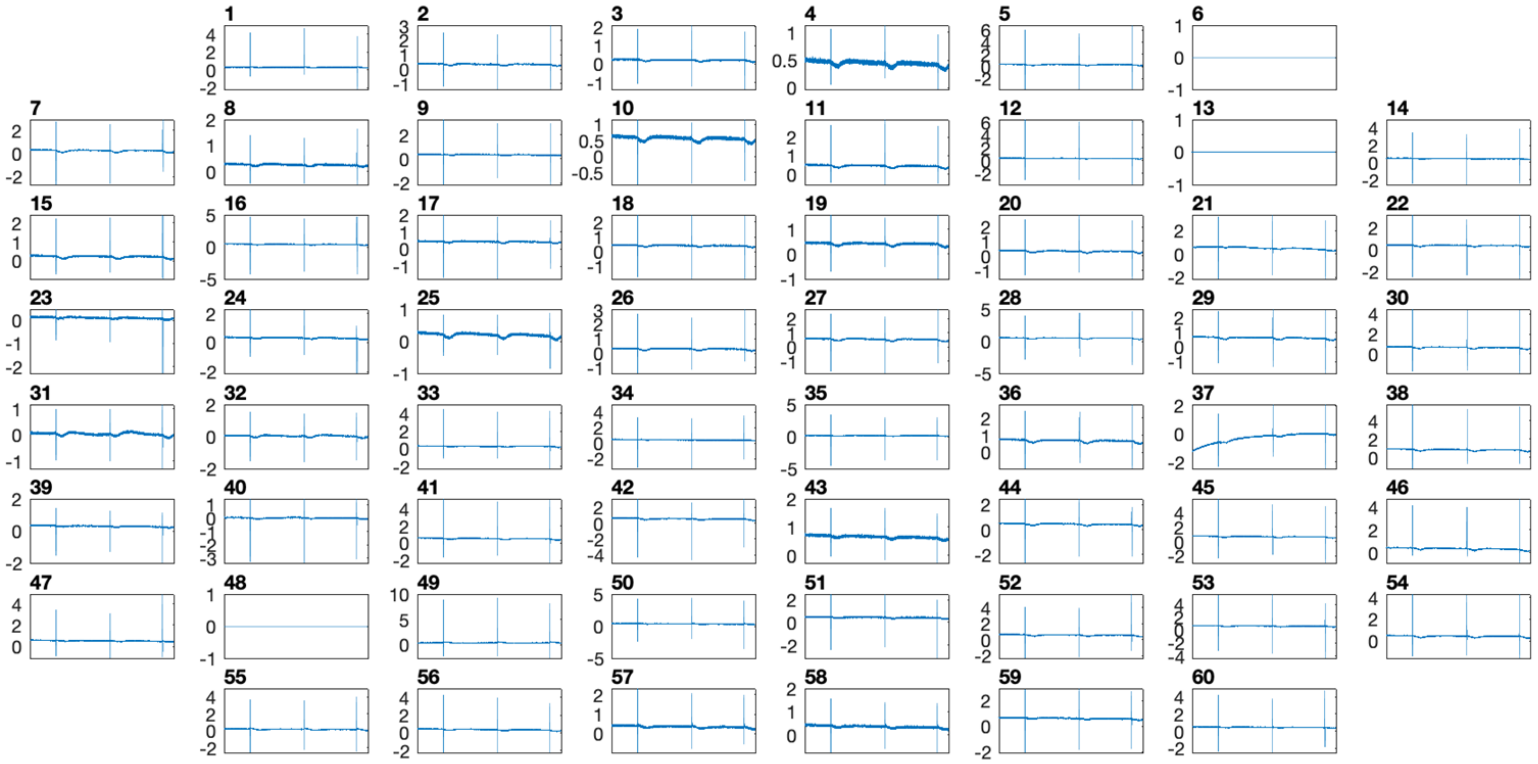

**Figure S7.** Multichannel action potential recording Before any electroporation (t=0-10s), all channels show extracellular signals except channel 6, which has a defective pin, and channels 13 and 48, which are large metal pads designed for pacing (not NEAs)

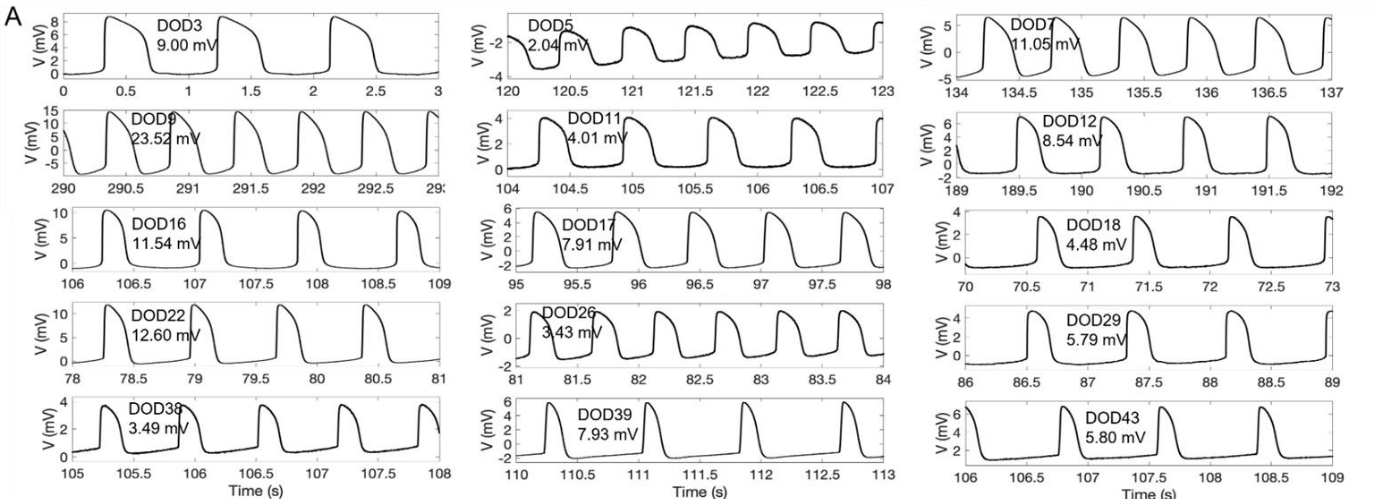


**Figure S8. Repeated recordings from the same cell over a month.** The same cell was measured 15 times from 3 days on device (DOD3) to DOD43.

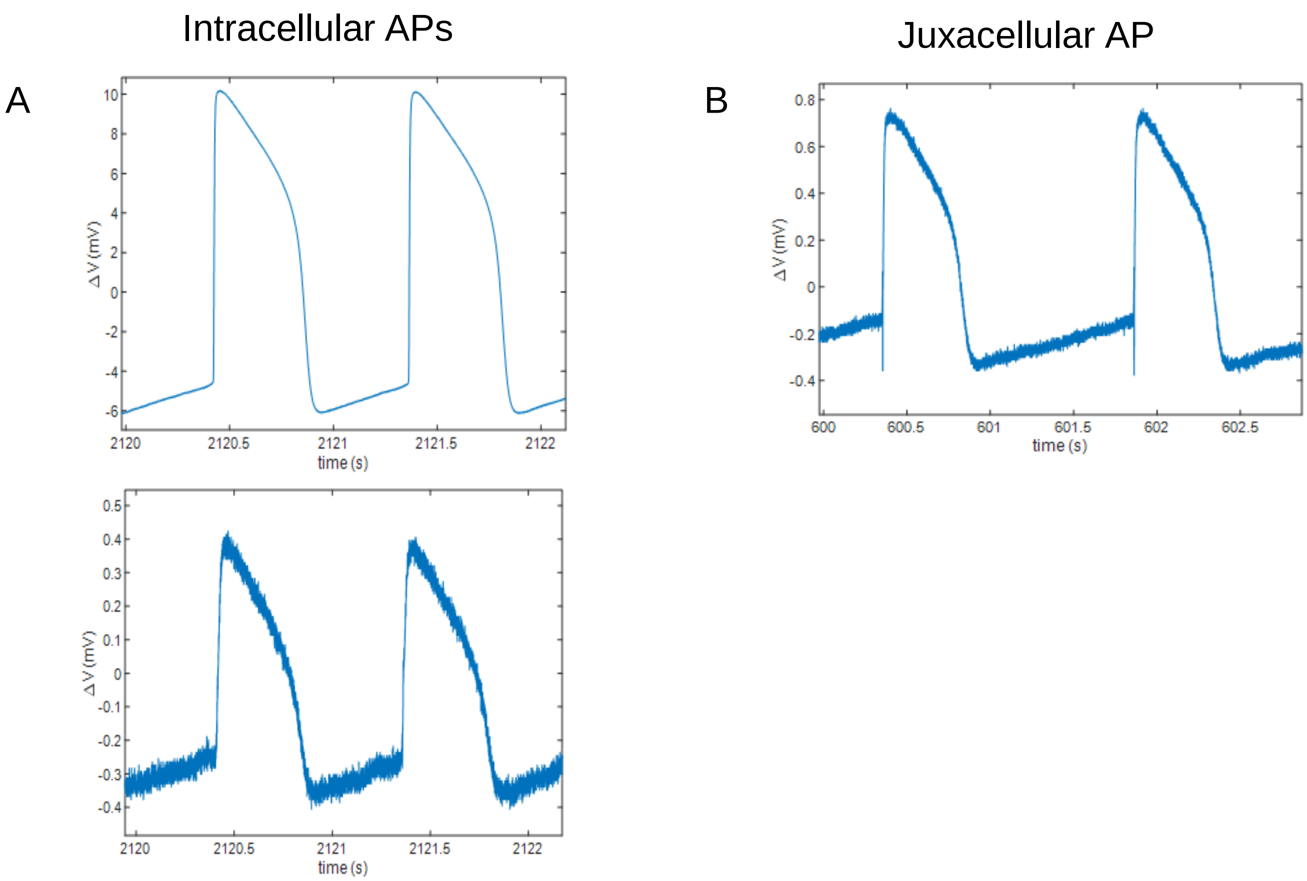


**Figure S9.** A) Examples of recordings with high or low amplitudes that are identified as intracellular APs in our analysis. B) Juxacellular (or hybrid) signal with intra and extracellular features - these are **not** counted as intracellular signals in our analysis.

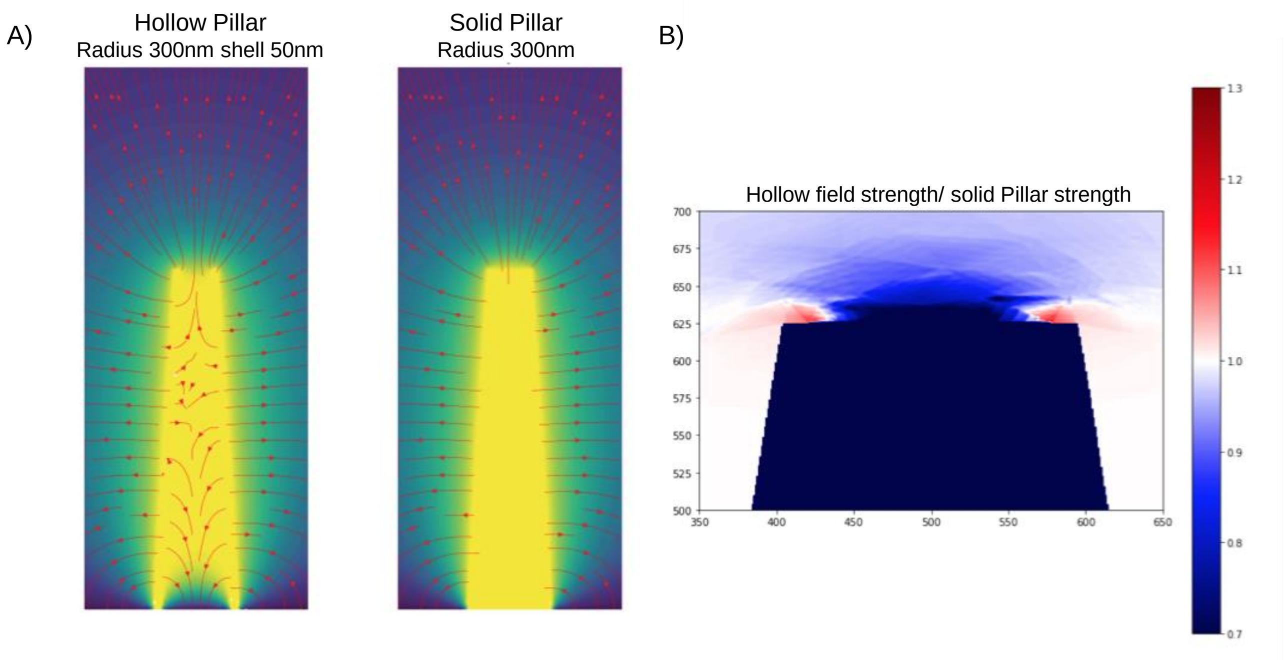


**Figure S10. Models of electric fields around hollow and solid nanopillars.** A) Hollow and full nanopillars are displayed with the voltage field overlaid in color (dark is V=0, bright is V=1). Some electric field lines are shown in red. The hollow pillar has field lines in its core as the voltage is not constrained there. B) The gradient of the voltage is computed for the full and hollow nanopillars to obtain the electric field at every spatial point. By computing the electric field magnitude and interpolating it spatially, the ratio of the field strengths can be compared at the same location. The color-code shows relative field strengths of the hollow nanopillar compared to the full one. An amplification of 30% (1.3) is observed for the hollow nanopillars.

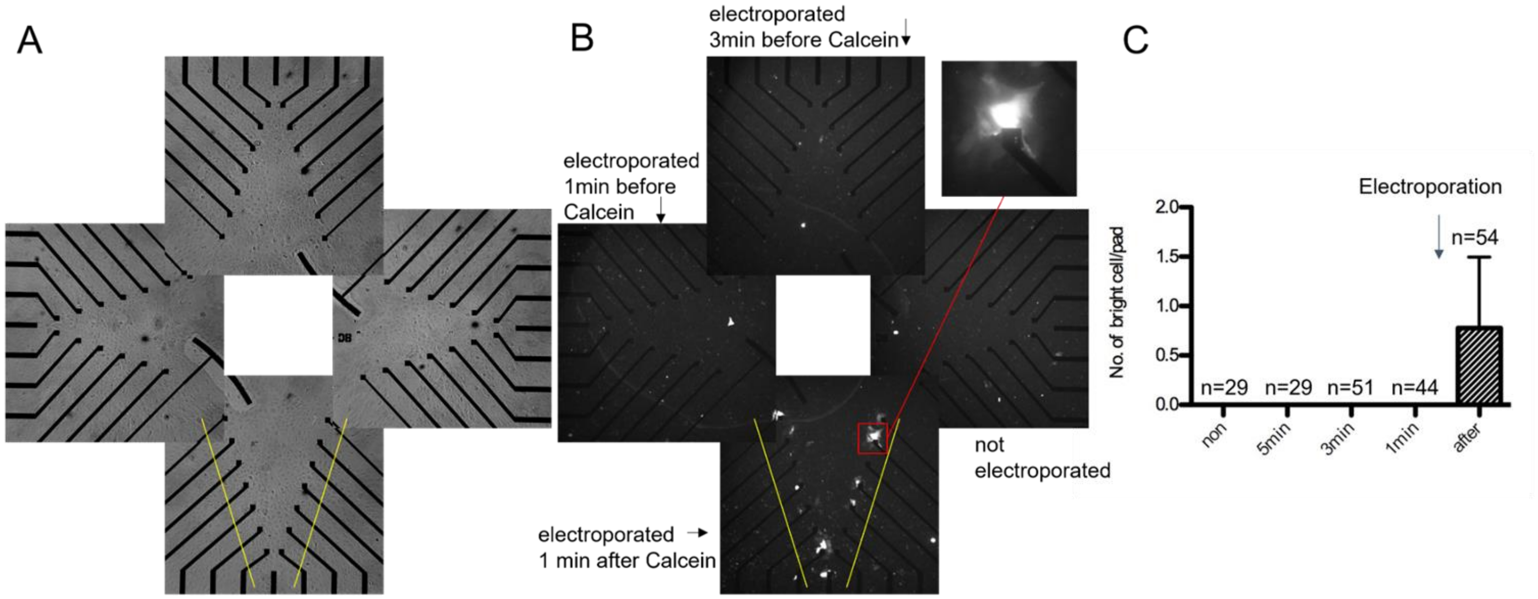


**Figure S11. Calcein entrance test of cell membrane permeability after NEA electroporation.** Each quadrant of the device was electroporated at 3 min before, 1 min before, or 1 min after adding 5μM Calcein, or not electroporated. Free Calcein was washed away 3 min after the last electroporation. **A**) Bright-field image. **B**) calcein imaging of the same view as the bright-field. **C**) Quantification of the number of bright cell per pad on each quadrant. Data comes from 4 independent experiments, in two among which 5 min electroporation before adding calcein was tested instead of non-poration. Tested pad number (n) is labeled on the bar graph. (mean ± SD).

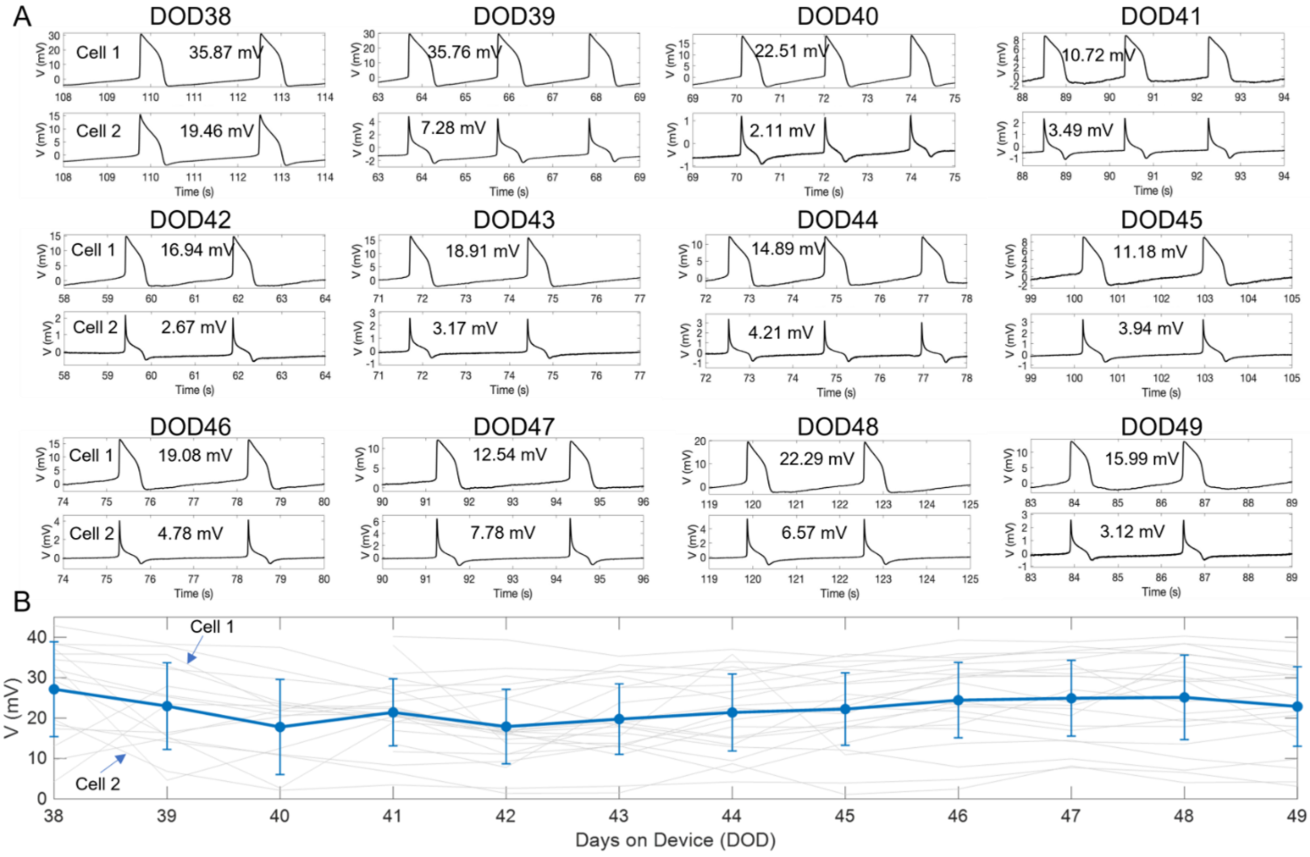


**Figure S12. Daily recording traces of two example cells in the same culture. A**) Example traces of two cells on the same device over 12 days. The two cells beat synchronously, but their iAPs waveforms differ. The waveform of cell 2 is rare, but the electrode consistently detects the same waveform over days. **B**) The plot of maximum iAP amplitude for each trace vs. DOD. The trace corresponding to the two cells in (A) is pointed out. The blue trace is the average of all cells over 12 days, which shows that the average iAP amplitude is stable during daily recordings.

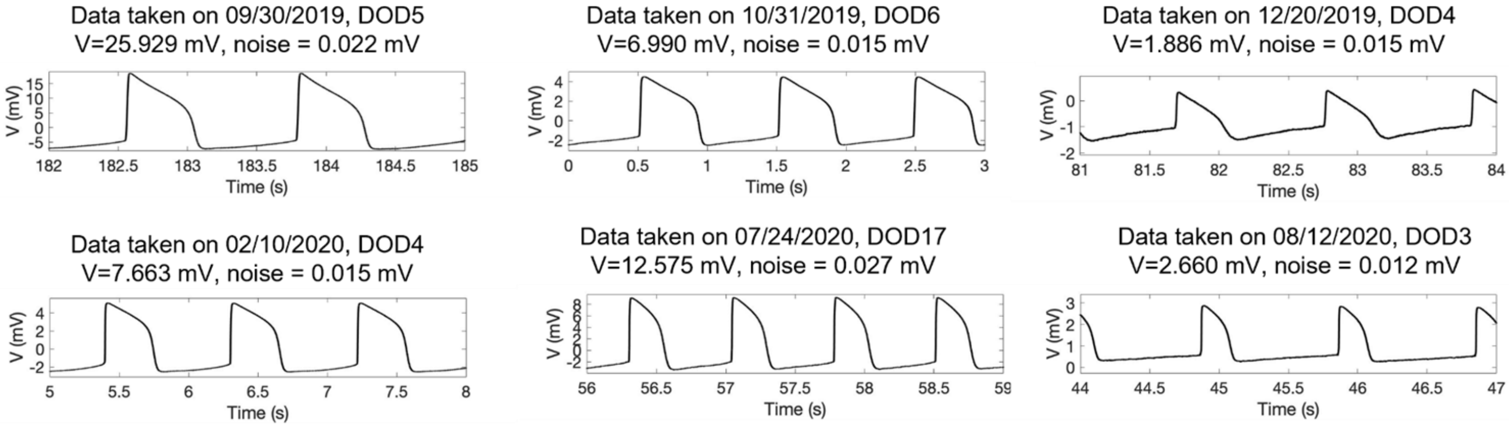


**Figure S13. The same device was repeatedly cleaned and re-used for new cell cultures.** The device maintains high signal quality and similar noise level when it has been re-used for many cultures over a year. Each plot is an example trace of a different culture.

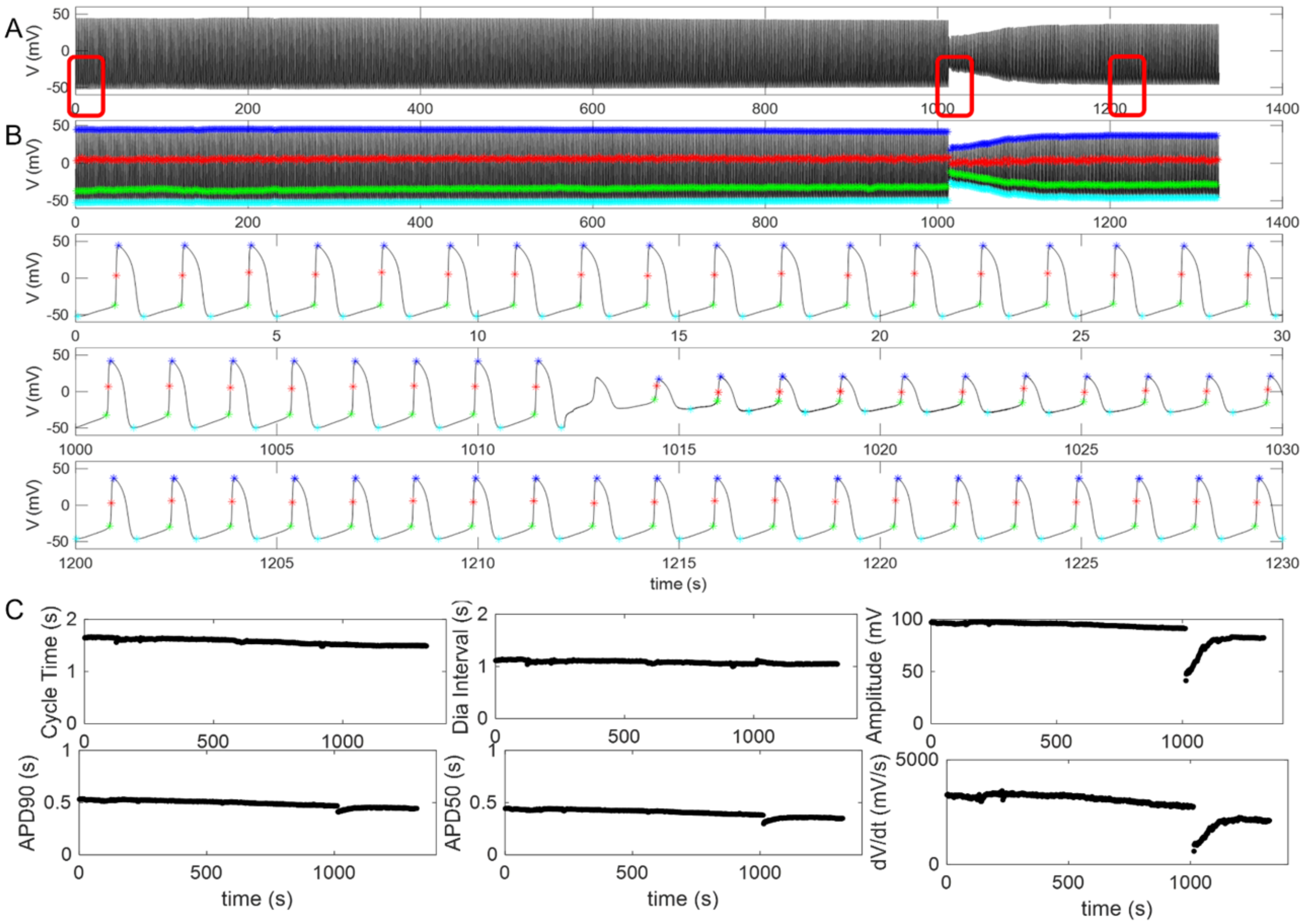


**Figure S14. A typical patch clamp recording trace.** A) A 1300-s long trace shows a sudden decrease of amplitude at 1012 second. B) Automated analysis of the trace that identifies the spiking point (red stars), the starting point (green stars), maximum point (bue stars), and the minimum point (cyan stars) of each iAP. The zoom-in windows show detailed waveforms. C) Time-dependent changes of the cycle time, diastolic interval, amplitude, APD90, APD50, and spiking velocity dV/dt. .

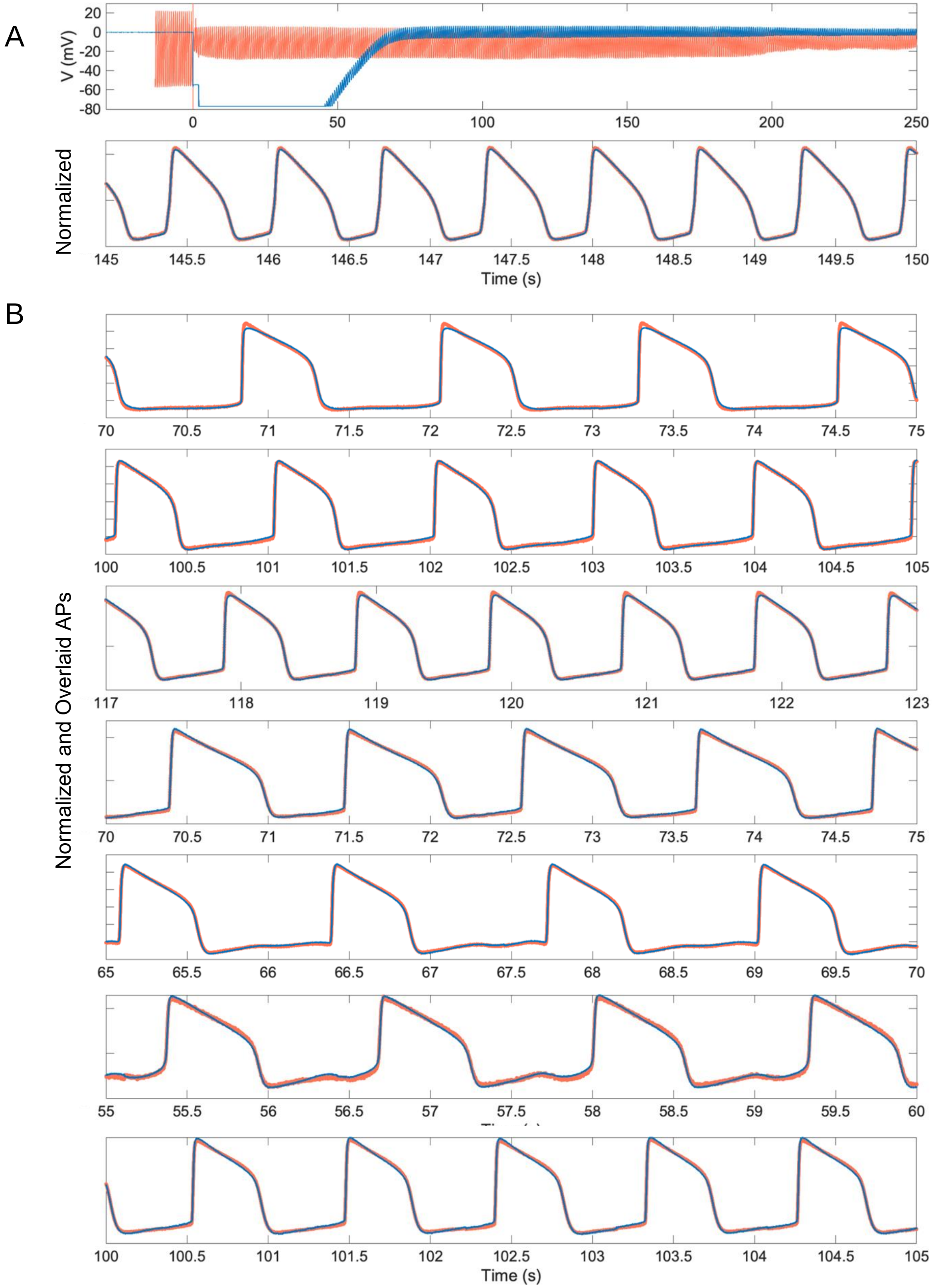


**Figure S15. Simultaneous Patch and NEA recordings.** A) Alignment of recordings by simultaneous patch and NEA recorded traces. The 5-second zoom-in window shows that the two traces overlap after amplitude scaling. B) 5-second zoom-in windows show the alignment of recordings by simultaneous Patch a and VNEA for 7 cells with distinct waveforms.

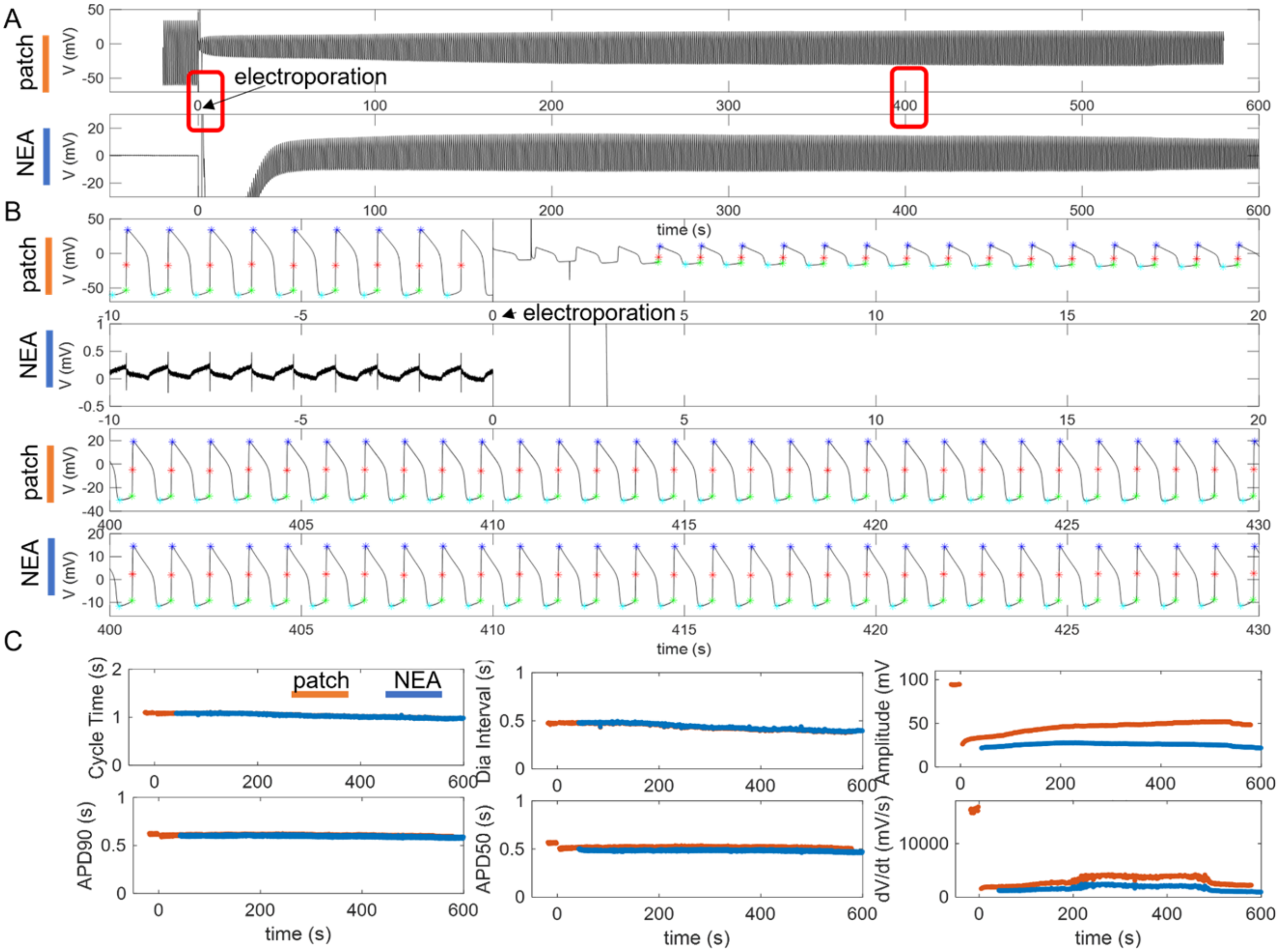


**Figure S16. Detailed analysis of the patch/NEA traces shown in Figure 3C.** A) Full length traces by patch and NEA. B) Zoom-in windows and automated analysis of the spiking point (red stars), the starting point (green stars), maximum point (bue stars), and the minimum point (cyan stars) of each iAP. C) Time-dependent changes of the cycle time, diastolic interval, amplitude, APD90, APD50, and spiking velocity dV/dt. .

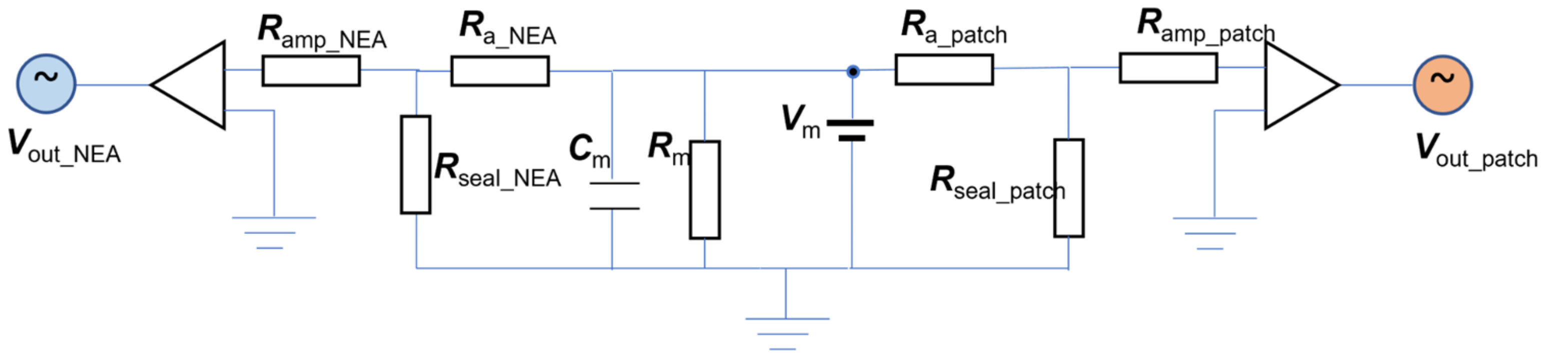


**Figure S17. Simplified patch and NEA simultaneous recording from the same cell.** This circuit diagram is to understand the low frequency signals. Double-layer capacitance is ignored. In the circuit diagram, $R_m$ and $C_m$ are the resistance and the capacitance of the cell membrane. $R_a$ is the access resistance and $R_{seal}$ is the sealing resistance. $R_{amp\_NEA}$ and $R_{amp\_patch}$ are the amplifier input resistances for the NEA and the patch amplifier. $V_m$ is the membrane voltage of the cell, and $V_{out}$ is the output voltage from the amplifier. The $V_{out\_patch}$ and $V_{out\_NEA}$ are linearly related to the biological signal $V_m$ through relations: $V_{out\_patch} = V_m * R_{seal\_patch} /(R_{seal\_patch} + R_{a\_patch})$ and $V_{out\_NEA} = V_m * R_{seal\_NEA} /(R_{seal\_NEA} +R_{a\_NEA})$. Before electroporation, the NEA access resistance $R_{a\_NEA}$ is extraordinarily large so NEA mostly records extracellular signal spikes that are dominated by capacitance coupling. The electroporation pulse locally permeabilizes the cell membrane around NEAs that drastically reduces $R_{a\_NEA}$ to achieve intracellular access. At the same time, through small capacitance coupling, the electroporation pulse can change the patch sealing and reduce $R_{seal\_patch}$. Because both $V_{out\_patch}$ and $V_{out\_NEA}$ are linearly related to the biological sigal $V_m$, they report the same iAP waveform despite different amplitudes.

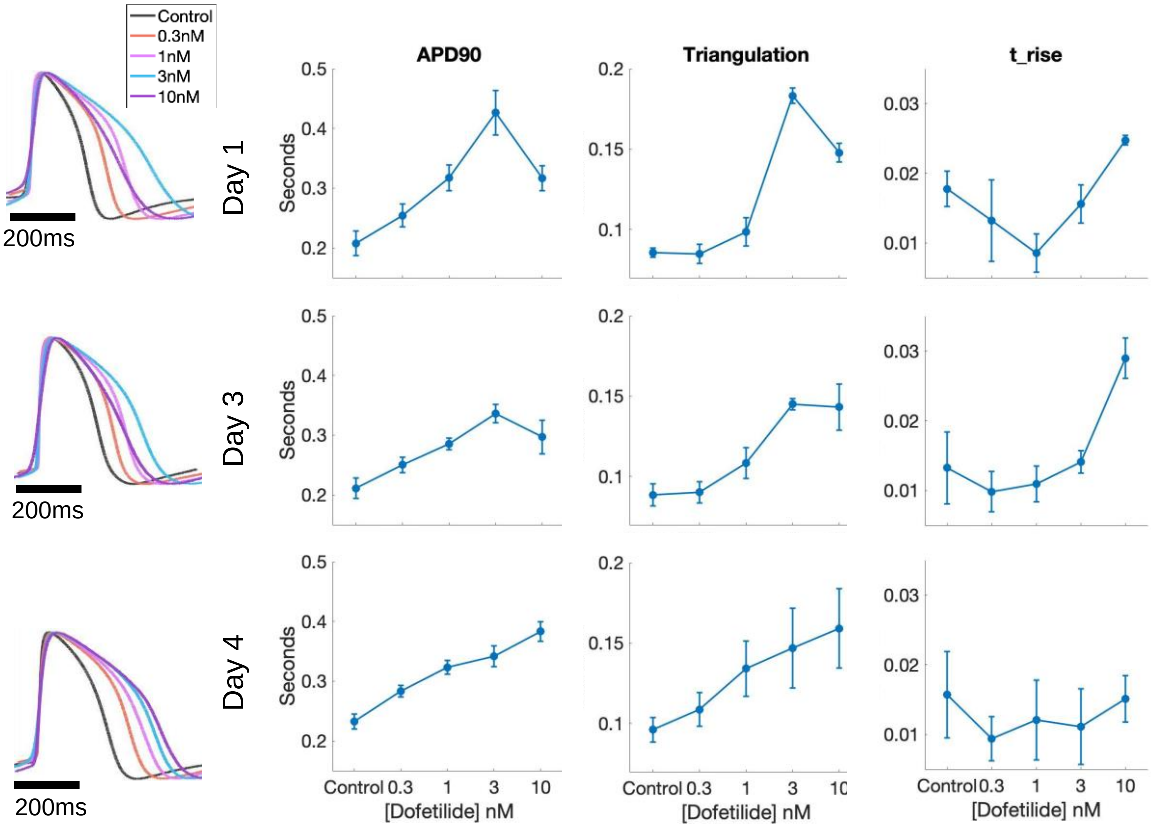


**Figure S18.** Independent pharmacology experiments on the same semi-hollow NEA device and the same cell culture over multiple days (Day 1-Day 4).
